# Remodeling of oxygen-transporting tracheoles drives intestinal regeneration and tumorigenesis

**DOI:** 10.1101/2021.01.23.427889

**Authors:** Vasilia Tamamouna, M. Mahidur Rahman, Monika Petersson, Irini Charalambous, Kristina Kux, Hannah Mainor, Verena Bolender, Buse Isbilir, Bruce A. Edgar, Chrysoula Pitsouli

**Author notes:** Equal-contributing authors (Vasilia Tamamouna, M. Mahidur Rahman).

## Abstract

The *Drosophila* tracheal system, as the functional equivalent of mammalian blood vessels, responds to hypoxia and transports oxygen throughout the body. Although the signaling pathways involved in tracheal development and the hypoxic response are well-studied, how adult tracheae interact with their target tissues is largely unknown. Here we show that the tracheae that serve the adult intestine are dynamic and respond to enteric infection, oxidative agents, and the development of gut tumors with increased terminal branching. Increased tracheation is necessary for efficient damage-induced intestinal stem cell (ISC)-mediated midgut regeneration and sufficient to drive ISC proliferation in the absence of damage. Gut damage or tumors induce Hif-1α/Sima, which, in turn, stimulates tracheole branching via the FGF(Brachless/Bnl)/FGFR(Breathless/Btl) signaling cascade. Bnl/Btl signaling is required both in the intestinal epithelium and the tracheal system for efficient damage-induced tracheal remodeling and ISC proliferation. We show that chemical or *Pseudomonas-*generated ROS directly affect the trachea and are necessary for branching and intestinal regeneration. Similarly, tracheole branching and the resulting increase in oxygen supply are essential for tumor growth in the midgut. Thus, we have identified a novel mechanism of visceral tracheal-intestinal tissue communication, whereby oxidative damage and tumors induce neo-tracheogenesis in adult *Drosophila.* This process is reminiscent of cancer-induced neo-angiogenesis in mammals.

## Introduction

The *Drosophila* tracheal system is an extended network of ramified gas-transporting tubes that transfer oxygen to all tissues. It is, thus, considered the functional equivalent of the mammalian vascular system ^1, 2^. The FGFR/Breathless (Btl) - FGF/Branchless (Bnl) receptor tyrosine kinase pathway is reiteratively utilized to direct tracheal formation during all developmental stages ^3–9^. The tracheal system encompasses specialized cells with exceptional plasticity. The terminal tracheal cells (TTCs) extend cytoplasmic tubular processes, the terminal branches (TBs) or tracheoles, towards tissues that need oxygen and are directly involved in gas exchange. TTCs form in response to developmental cues during development, but oxygen levels regulate TB remodeling. Low oxygen (hypoxia) induces TTCs to extend more TBs, whereas increased oxygen (hyperoxia) suppresses TB formation ^10–12^. Therefore, as oxygen sensors, the TTCs are equivalent to the tip cells of the mammalian vasculature ^13^, which are involved in sprouting angiogenesis.

Hypoxia induces the expression of the chemoattractant FGF/Bnl in non-tracheal tissues, and this directs the newly formed TBs of TTCs that express the FGFR/Btl on their surface ^1, 14^. The formation of new TBs depends on the Hypoxia-Inducible Factor 1α (Hif-1α)/Similar (Sima), and the Hif-propyl hydroxylase (Hph)/Fatiga (Fga), the latter acting as an oxygen sensor ^11^. Upon hypoxia, Sima is expressed in both tracheal cells and non-tracheal cells to induce *FGFR/btl* and *FGF/bnl*, respectively, and it is necessary for TB remodeling in *Drosophila* larvae ^11^. Thus, *Drosophila* TTC remodeling is controlled by the Hif-1α pathway via the activation of the FGFR-FGF pathway, similar to angiogenesis in mammals ^15, 16^. Although the embryonic development of the *Drosophila* trachea is well-characterized, our knowledge about the formation, maintenance and remodeling of the tracheal system in the adult fly is limited.

The *Drosophila* adult midgut has emerged as an excellent system to study stem cell regeneration in response to chemical stress, infection, aging, as well as tissue communication ^17–20^. The endoderm-derived midgut is an apicobasally polarized epithelium comprised of intestinal stem cells (ISCs), and their progeny, the transient enteroblasts (EBs) and pre-enteroendocrine cells (pre-EEs), which differentiate to lumen-facing absorptive enterocytes (ECs) and secretory enteroendocrine cells (EEs), respectively ^21–23^. A layer of visceral muscle (VM) composed of longitudinal and circular fibers ensheaths the epithelium. The VM supports midgut peristalsis and also supplies a basement membrane and signaling molecules that are essential components of the ISC niche ^24, 25^. Outside the VM lie the visceral tracheae that branch extensively and penetrate the VM to access the intestinal epithelium ^26^ to supply it with oxygen internalized via the spiracles ^27, 28^. The intestinal tracheae are known to be dynamic and to respond to nutritional cues ^29^, have been proposed to contribute niche signals for ISC maintenance ^26^, and have been shown to associate with metastatic tumors ^30^. However, a mechanistic study on the role of the intestinal trachea in gut regeneration and tumorigenesis has not yet been described.

Bacterial pathogens and damage-inducing chemicals activate conserved signaling pathways that promote regeneration of the damaged intestine. For example, oral infection with the pathogenic bacteria *Pseudomonas entomophila* (*P.e.*) or *Pseudomonas aeruginosa* (*P.a*.) damages the intestinal epithelium and induces ISC-mitoses, which facilitate regeneration ^31–33^. Pyocyanin, a ROS-generating secreted virulence factor of the pathogenic *P.a.* strain PA14, is sufficient to induce intestinal regeneration ^33^. Ingestion of other ROS-like chemicals (hydrogen peroxide (H_2_O_2_), Paraquat (PQ)), and tissue-damaging agents (bleomycin, dextran sulfate sodium (DSS)) also lead to ISC-mediated gut regeneration ^34–37^. Regeneration is achieved via the induction of paracrine signals from the ISC niche that activate conserved cell signaling pathways in the gut epithelium, including the JNK, Jak/Stat, Wnt/Wingless (Wg), Hippo/Yki and EGFR ^18, 31, 34, 38–43^.

Oxygen is essential for tissue metabolism. Abundant oxygen, and oxygen-derived ROS play essential roles in cellular respiration and oxidative damage, and often act as signaling second messengers ^44^. Oxygen can increase mitochondrial respiratory chain activity resulting in elevated O_2_, OH or H_2_O_2_, which, in turn, may cause macromolecular cellular damage. In addition, cell death-induced ROS activate the JNK, p38^MAPK^ and Jak/Stat pathways to promote regeneration ^45, 46^. Since ROS can be both damaging and essential for regeneration, tight control of their levels is necessary for tissue homeostasis. ROS homeostasis is controlled by antioxidant enzymes, such as superoxide dismutase, catalase, and glutathione S-transferase ^47^. Tumor cells exhibit high levels of both growth-promoting ROS and protective anti-oxidant pathways, including the Nrf2 pathway ^48, 49^. Although ROS signaling is involved in tissue injury response and tumor development, the mechanism, if any, of how ROS modulate tracheation remains unknown.

In this study, we have investigated the signals that mediate communication between the visceral trachea and the intestinal epithelium. We show that the adult midgut-associated tracheae increase their terminal branching in response to infection, oxidative stress, and tumor development. Increased TTC branching is, in turn, necessary for a robust ISC regenerative response, and tumor growth. By comparing responses in normoxia and hypoxia, we show that the availability of oxygen, which is transported to the epithelium via the TTCs, is essential for the ISC-mediated regeneration response to infection or oxidative stress.

## Results

### Intestinal tracheal remodeling induced by bacterial infection and oxidative damage is necessary for ISC proliferation

Previous studies have suggested that the *Drosophila* trachea can actively regulate the metabolic status of adult tissues ^26, 29^. To assess if different damage-causing and regeneration-promoting agents can induce intestinal tracheal branching, we subjected wild-type flies to different stresses (Fig. 1a-l). We labeled the trachea with fluorescent proteins either via the Gal4-UAS system ^50^ using the trachea-specific *btl-Gal4>UAS-srcGFP* reporter ^51^ or via the QF-QUAS system ^52^ using the *QF6>QUAS-mtdTomato* reporter that is expressed in the trachea (Fig. 1; Extended Data Fig. 1). The flies were either infected orally with pathogenic bacteria (*Pseudomonas aeruginosa* (*P.a.*), *Pseudomonas entomophila* (*P.e.*)) or fed with oxidative agents such as hydrogen peroxide (H_2_O_2_) or Paraquat (PQ). These stresses are known to induce ISC proliferation by damaging the intestinal epithelial cells ^31, 33, 35, 37^. As expected, all treatments led to a significant increase in *esg>GFP+* cells in the midgut (Extended Data Fig. 1a) and ISC proliferation (Fig. 1h,j,l, and Extended Data Fig. 1b). Strikingly, *Pseudomonas* infection and oxidative agents also induced an increase in the midgut-associated tracheal branches (Fig. 1a-f; Extended Data Fig. 1c-e). To quantify the midgut tracheal coverage, we used confocal images with fluorescently-labeled trachea or bright-field images of midgut preparations without detergent. In the latter case, the air was maintained in the tubes and offered contrast. Then, we applied the NeuronJ Plugin of the ImageJ software (NIH) on the images to measure the total length of the tracheal branches overlaying the midgut, which was normalized to the gut surface. In a few cases (e.g. Fig. 1k), we measured the total number of terminal branches in specific midgut regions. Importantly, quantification of the images confirmed the imaging observations (Fig. 1g,i,k; Extended Data Fig. 1c). To exclude the possibility that extensive tracheal branching was caused by increased TTC numbers, we measured the number of TTCs in uninfected and *P.a.*-infected midguts. We found that infection did not alter the number of midgut TTCs (Fig. 1m). Further, we did not observe any intestinal TTC undergoing mitosis in baseline conditions or under any treatment. Thus, our results indicate that direct epithelial damage caused by pathogenic bacteria or ROS increases ISC proliferation and also reorganizes the adult tracheoles surrounding the intestine.

**Figure 1.**
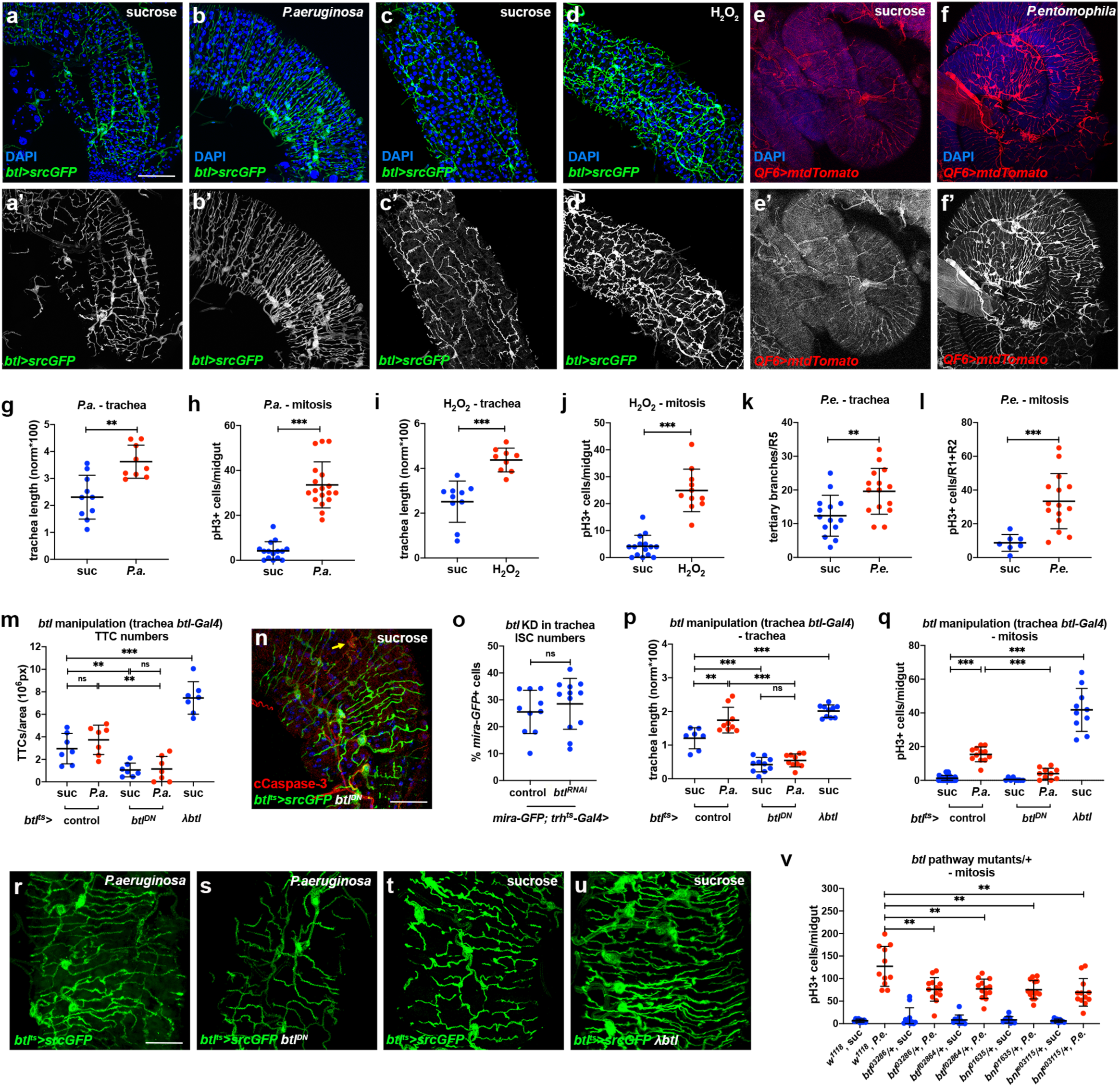
Remodeling of visceral TTCs covering the midgut is associated with ISC proliferation. **a-f.** The adult intestinal trachea is increased upon *P.a.* infection (a, b), administration of H_2_O_2_ (c, d) and *P.e.* infection (e, f). The intestinal TTCs were labeled with *btl-Gal4>UAS-srcGFP* in a-d and with *QF6>QUAS-mtdTomato* in e-f, and DAPI (blue) labeled all nuclei in a-f. The R5 midgut region was imaged in a-b, the R2 region in c-d and the R4-R5 region in e-f. Single channel images of the trachea are shown in a’-f’. **g-l.** Quantification of TTC branching and mitosis upon *P.a.* infection (g, h), administration of H_2_O_2_ (i, j) and *P.e.* infection (k, l). **m-** Quantification of TTCs upon trachea-specific *btl* manipulation with or without *P.a.* infection. **n**. Tracheae (green) and cleaved Caspase-3 (red) upon trachea-specific *btl* knockdown. **o.** Quantification of *mira-GFP*-positive ISCs upon trachea-specific *btl* knockdown via *trh-Gal4*. **p-q.** Quantification of TTC branching (p) and mitosis (q) upon trachea-specific *btl* manipulation with or without *P.a.* infection. **r-u**. Tracheae (green) of *P.a.*-infected R5 regions of the midgut in *btl-Gal4>UAS-srcGFP* control (r) and *btl-Gal4>UAS-srcGFP, UAS-btl^DN^*-expressing (s), as well as control (t) versus *btl-Gal4>UAS-srcGFP, UAS-λbtl*-expressing flies (u). **v.** Quantification of midgut mitosis of flies heterozygous for *btl* pathway mutations with or without *P. e.* infection compared to controls. Scale bars: 75 μm in a-d, 60 μm in e-f, 37.5 μm in n, r-u. Error bars correspond to standard deviation. Statistical significance (t-tested): ns, not significant, * 0.01<*p*≤0.05, ** 0.001<*p*≤0.01 and *** *p*≤0.001.

Since the FGFR/Breathless (Btl) pathway is integral to tracheal formation and remodeling during *Drosophila* development ^1, 2^, we reasoned that its manipulation might affect the dynamics of adult intestinal tracheae. To assess whether the FGFR receptor, Btl, is necessary in the trachea for maintenance and remodeling, we overexpressed a dominant-negative form of Btl, *btl^DN^*, specifically in the adult tracheal system. We noticed that adult files with trachea-specific *btl* inhibition under *btl^ts^-Gal4* exhibited reduced numbers of intestinal TTCs in both uninfected and *P.a.*-infected conditions compared to control flies (Fig. 1m). This TTC reduction was accompanied by TTC apoptosis evident by cleaved Caspase-3 staining (Fig. 1n). Importantly, trachea-specific *btl* knockdown did not alter the number of *mira-GFP+* ISCs (Fig. 1o) in uninfected midguts, thereby confirming that the mitosis impairment in *btl* knockdown flies was not due to reduced number of ISCs. Upon *P.a.* infection, flies expressing *btl^DN^* in the adult trachea under *btl^ts^-Gal4* exhibited reduced tracheal branching relative to controls (Fig. 1r-s). We confirmed these *btl^DN^* results with two additional trachea-specific drivers (*trh-Gal4* and *dSRF-Gal4*) (Extended Data Fig. 2a-b, h-i). In addition, we overexpressed a dimerized constitutively-active form of the Btl receptor, *λbtl*, specifically in the trachea and found that it was sufficient to increase intestinal TTCs and induce intestinal tracheal remodeling in the absence of infection (Fig. 1m, t-u; Extended Data Fig. 2c-d, j-k). Quantification of the total intestinal tracheal length normalized to the intestinal surface using NeuronJ confirmed the observations derived from the images (Fig. 1p; Extended Data Fig. 2f, m). Therefore, trachea-specific increase and decrease of *btl* activity leads to increased and decreased tracheole density, respectively. To test whether intestinal tracheal remodeling is necessary for the regenerative response of the ISCs to damage, we assessed ISC proliferation in flies with trachea-specific manipulation of *btl*. We found that flies expressing *btl^DN^* in the trachea exhibited less mitosis upon *P.a.* infection, whereas flies expressing *λbtl* in the trachea exhibited increased mitosis in the absence of damage (Fig. 1q; Extended Data Fig. 2g, n). Furthermore, to underscore the necessity for the FGFR/Btl pathway in intestinal regeneration upon infection, we subjected the *btl^MI03286^*, *btl^f02864^, bnl^MI01635^* and *bnl^e03115^* mutants as heterozygotes to *P.e.* infection and found a significant decrease in their ability to regenerate the intestine (Fig. 1v). Collectively, these results indicate that FGFR/Btl receptor-mediated tracheal branching is critical for effective ISC regeneration.

### The FGF/FGFR signaling is necessary in both the epithelial cells and the tracheae for effective tracheal remodeling and ISC proliferation in response to damage

The *Drosophila* FGF homolog, Branchless (Bnl), is essential for the directional migration of terminal tracheal cells during embryonic development, as well as in larvae in response to hypoxia ^10, 11, 53^. To test whether Bnl is involved in intestinal tracheal remodeling in response to damage, we examined *bnl* induction in response to different damage regimes. Using *bnl-Gal4>UAS-srcGFP* as a reporter, we monitored the expression of *bnl* in the adult midgut. We found that *bnl* was expressed in a few epithelial cells of the midgut and that the number of *bnl*-expressing cells was robustly increased when the flies were fed with *P.a.*, H_2_O_2,_ or PQ (Fig. 2a-d; Extended Data Fig. 3). To confirm that *bnl* is induced at the mRNA transcript level, we performed RT-qPCR analysis in whole midguts of wild-type *w*^1118^ flies at the same time points as those used for reporter imaging. We found that *bnl* mRNA was induced by more than 1.5-fold by *P.a.* infection and PQ, but not H_2_O_2_. The lack of *bnl* mRNA induction by H_2_O_2_ in whole midgut tissue is possibly due to the regional effect of H_2_O_2_ as seen on the *bnl* reporter (Extended Data Fig. 3), which can affect the RT-qPCR sensitivity (Fig. 2e). To identify the specific cell types where *bnl* is expressed, we performed co-localization experiments of *bnl-Gal4>UAS-srcGFP* with the cell type specific markers *esg-lacZ*, *Dl-lacZ*, *Su(H)-lacZ*, and Prospero, which report midgut progenitors (ISCs and EBs), ISCs, EBs and EEs, respectively. We also identified differentiated ECs by their large size. We found that *bnl* was expressed and induced in all the epithelial midgut cell types. We did not detect any staining of the reporter outside the epithelial cell layer, such as in the VM or trachea (Fig. 2a-d and data not shown). To assess whether Bnl is involved in midgut tracheal remodeling, we knocked-down *bnl* using tissue-specific RNAi in adult intestinal progenitor cells (ISCs and EBs) or ECs using *esg^ts^-Gal4* or *Myo1A^ts^-Gal4*, respectively. In both cases, *bnl* knockdown led to reduced tracheal coverage in *P.a.* or *P.e.*-infected midguts (Fig. 2f,j; Extended Data Fig. 4a,d). In addition, the ISC proliferation rate in response to either *P.a.* infection (Fig. 2g-k) or *P.e.* infection (Fig. Extended Data Fig. 2c,f) was reduced, suggesting that *bnl* is necessary for an effective regenerative response. To test whether *bnl* overexpression is sufficient to induce tracheal remodeling in the absence of damage, we overexpressed *bnl* in either intestinal progenitors (ISCs and EBs) or in ECs. Not surprisingly, *bnl* overexpression was sufficient to induce tracheal remodeling (Fig. 2h-l). Remarkably, *bnl* overexpression in intestinal progenitors or ECs was also sufficient to induce ISC mitoses (Fig. 2i,m). Importantly, *bnl* knock-down in intestinal progenitors did not affect the viability and the total numbers of progenitor cells (Fig. 2n-q; Extended Data Fig. 4g). Collectively, our data suggest that damage-induced *bnl* expression in the midgut epithelial cells is necessary for ISC proliferation, and that *bnl* is sufficient for TTC remodeling and induction of ISC proliferation in the absence of damage.

**Figure 2.**
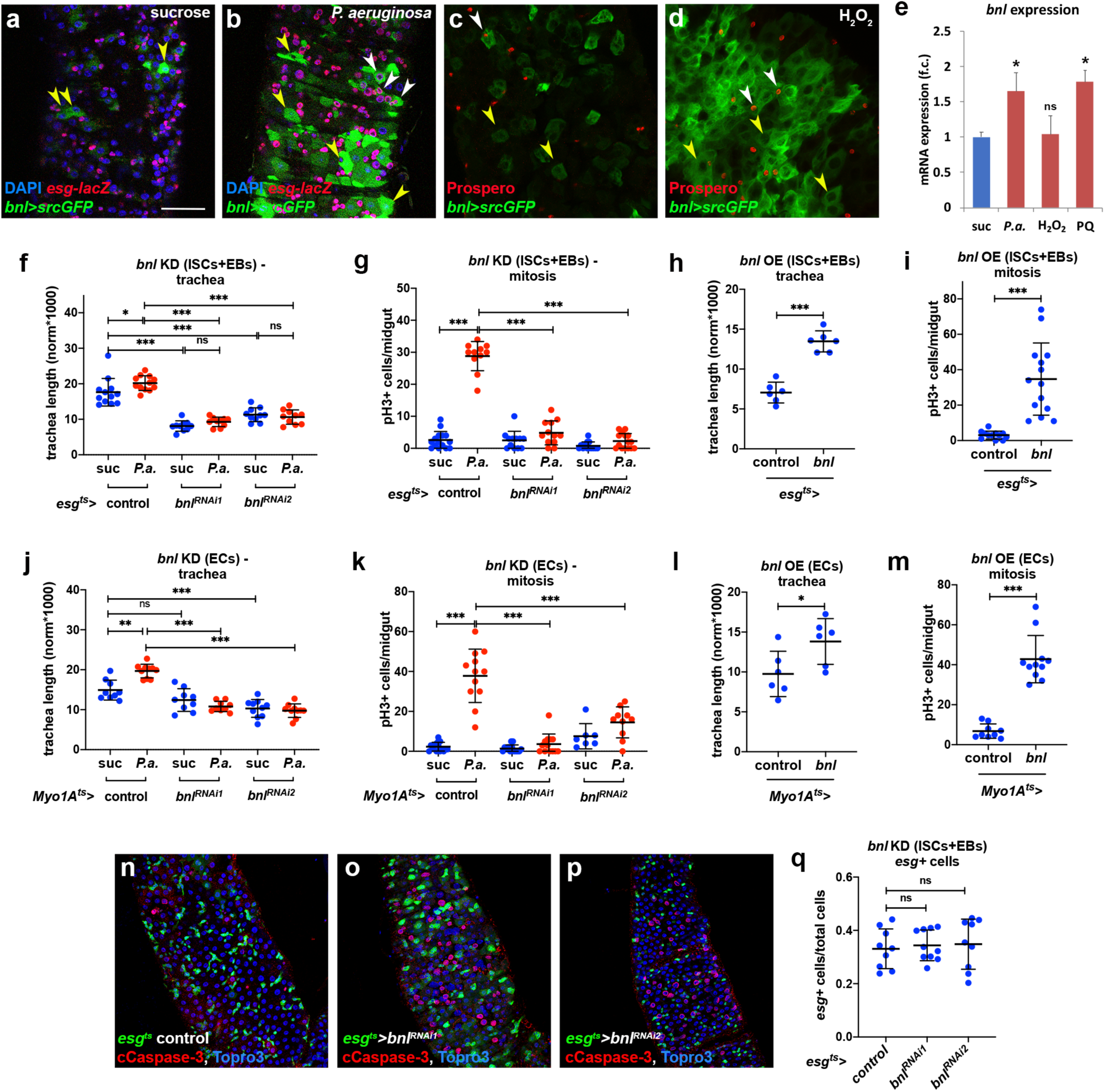
The FGF/Bnl is expressed in the midgut epithelium and controls ISC proliferation and TTC remodeling. **a-b.** Control (a) and *P.a.*-infected (b) *bnl-Gal4>UAS-srcGFP* midguts (R5 region) with *esg-lacZ*-labeled (red) ISCs and EBs, and DAPI (blue) staining all the nuclei. **c-d.** Control (c) and H_2_O_2_-fed (d) *bnl-Gal4>UAS-srcGFP* midguts (R2 region) with Prospero-labeled (red) EEs, and DAPI (blue) staining all the nuclei. Yellow arrowheads indicate *bnl*-positive ECs (big cells) and white arrowheads indicate *bnl*-positive *esg*-positive progenitors (a-b) and EEs (c-d). **e**. RT-qPCR for *bnl* mRNA expression in *w*^1118^ whole adult midguts upon different stresses. The average and standard deviation of biological triplicate experiments are plotted. **f-g**. Quantification of TTC branching (f) and mitosis (g) upon progenitor-specific *bnl* knockdown with or without *P.a* infection. **h-i**. Quantification of TTC branching (h) and mitosis (i) upon progenitor-specific *bnl* overexpression. **j-k.** Quantification of TTC branching (j) and mitosis (k) upon EC-specific *bnl* knockdown with or without *P.a* infection. **l-m**. Quantification of TTC branching (l) and mitosis (m) upon EC-specific *bnl* overexpression. **n-p**. Midguts (R5 region) of control (n) and *esg^ts^-Gal4* progenitor-specific *bnl* knockdown (o-p), stained for ISCs+EBs (green), cleaved Caspase-3 (red) and Topro-3 (blue). **q**. Quantification of *esg*+ ISCs+EBs in control and progenitor-specific *bnl* knockdown. Scale bars: 37.5 μm in a-d, 75 μm in n-p. Error bars correspond to standard deviation. Statistical significance (t-tested): ns, not significant, * 0.01<*p*≤0.05, ** 0.001<*p*≤0.01 and *** *p*≤0.001.

We did not observe *btl* outside the trachea or *bnl* outside the intestinal epithelium (Fig. 1a-d, Fig. 2a-d). Nevertheless, and because the reporters might not precisely highlight all the cells expressing the genes of interest, we decided to test *btl* and *bnl* requirements in the intestinal epithelium and the trachea, respectively. To assess if *btl* is necessary in the intestinal progenitors (ISCs and EBs) or ECs, we inactivated *btl* by overexpressing *btl^DN^* or *btl^RNAi^* under *esg^ts^-Gal4* and *Myo1A^ts^-Gal4*, respectively. We found that both tracheal branching and ISC mitosis were impaired when *btl* was inhibited in the epithelium upon *P.a.* or *P.e.* infection (Fig. 3a-d; Extended Data Fig. 4a-b,d-e). Importantly, progenitor-specific downregulation of *btl* did not affect the number of intestinal *esg+* progenitors (Extended Data Fig. 4g-h). Thus, *btl* is necessary in the intestinal epithelial cells for mitosis and tracheal remodeling upon infection.

**Figure 3.**
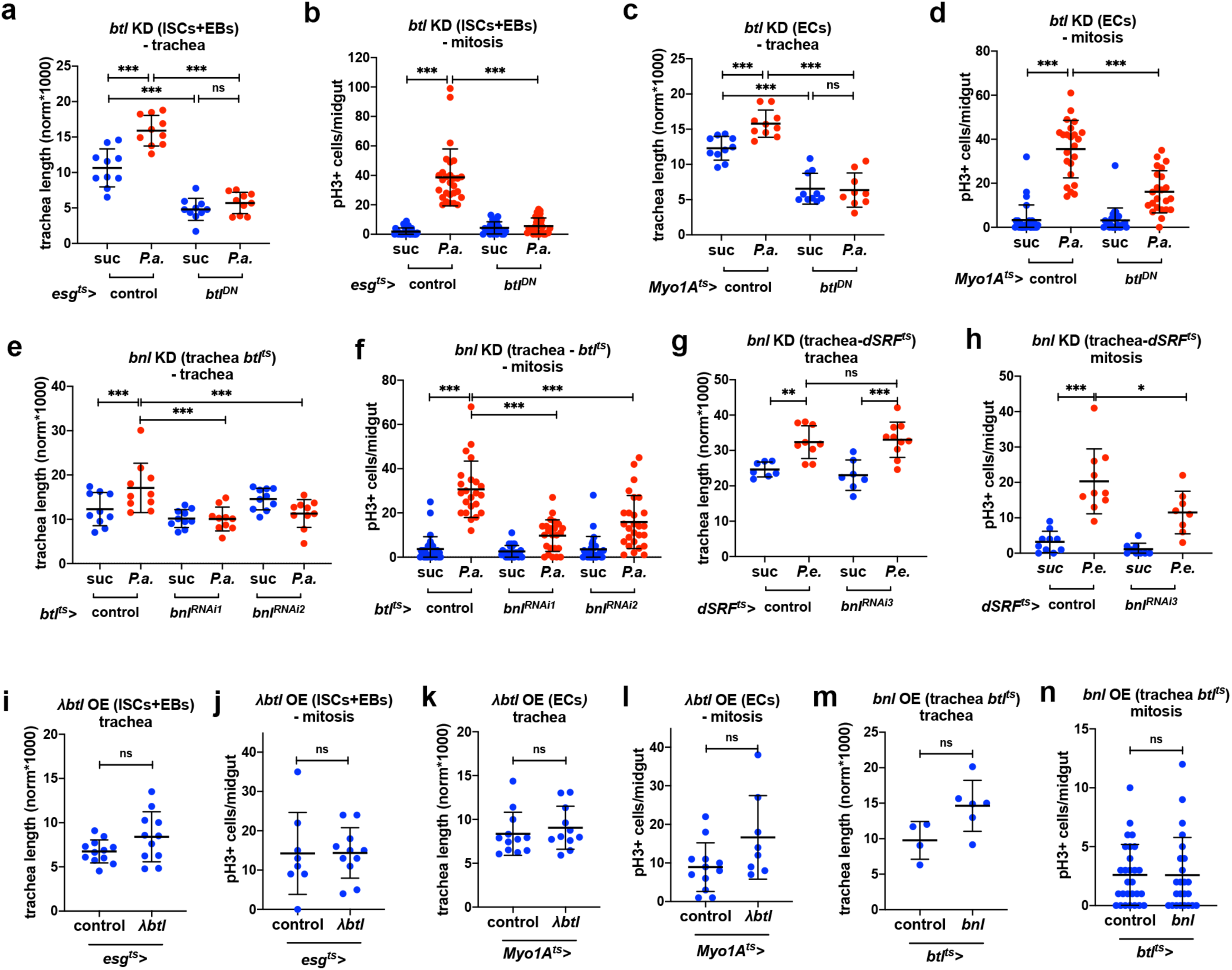
Damage-induced TTC remodeling and ISC mitosis require epithelial FGFR/Btl and tracheal FGF/Bnl. **a-d**. Quantification of TTC branching (a,c) and mitosis (b,d) upon progenitor-specific (a-b) and EC-specific (c-d) *btl* knockdown with or without *P.a.* infection. **e-f**. Quantification of TTC branching (e) and mitosis (f) upon trachea-specific *btl* knockdown via *btl^ts^-Gal4* with or without *P.a.* infection. **g-h**. Quantification of TTC branching (g) and mitosis (h) upon trachea-specific *btl* knockdown via *dSRF^ts^-Gal4* with or without *P.e.* infection. **i-j**. Quantification of TTC branching (i) and mitosis (j) upon progenitor-specific *λbtl* overexpression. **k-l**. Quantification of TTC branching (k) and mitosis (l) upon EC-specific *λbtl* overexpression. **m-n**. Quantification of TTC branching (m) and mitosis (n) upon trachea-specific *λbtl* overexpression via *btl^ts^-Gal4*. The flies were uninfected in i-n. Error bars correspond to standard deviation. Statistical significance (t-tested): ns, not significant, * 0.01<*p*≤0.05, ** 0.001<*p*≤0.01 and *** *p*≤0.001.

To assess if *bnl* functions in the tracheal cells, we knocked down its expression in the adult trachea via the *btl^ts^-Gal4* using two independent RNAi lines. We found that *bnl* is necessary in the adult trachea for both damage-induced tracheal branching and mitosis (Fig. 3e-f). Interestingly, *bnl* knock-down in the adult trachea using the *dSRF^ts^-Gal4* also showed that *bnl* is required in the tracheal cells for ISC mitosis upon infection, but tracheal remodeling was not affected in this genotype (Fig. 3g-h). Thus, *bnl* is necessary in the intestinal trachea for mitosis and damage-induced tracheole remodeling.

To test if epithelial *btl* is sufficient to induce ISC mitosis and visceral trachea remodeling, we overexpressed activated *btl* in progenitor cells or ECs using the *esg^ts^-Gal4* and *Myo1A^ts^-Gal4*. Neither progenitor- nor EC-specific *λbtl* expression affected tracheal remodeling and ISC mitosis (Fig. 3i-l). To assess if tracheal *bnl* is sufficient for the same phenotypes, we overexpressed *bnl* under *btl^ts^-Gal4*. We found that *bnl* is not sufficient to induce mitosis or tracheal remodeling when overexpressed in the trachea (Fig. 3m-n). Therefore, neither epithelial *btl* nor tracheal *bnl* are sufficient to induce mitosis and tracheal remodeling, indicating that other factors are also needed.

### The Hif-1*α* homolog Sima is required for tracheal remodeling and ISC proliferation

Sima, the *Drosophila* homolog of Hypoxia-inducible factor 1*α* (Hif-1*α*), controls Btl activity and tracheal remodeling in response to low oxygen during *Drosophila* larval development ^11^, and is induced in response to epithelial damage and tumors ^54, 55^. In mammalian cells, Hif-1*α* activity can also be controlled by ROS, produced by mitochondria in moderate hypoxia, and can inhibit the PHD/Hph proteins that promote Hif-1*α* degradation ^56–58^. In normoxia, Hif-1*α* activity can be controlled by exogenous ROS, e.g. H_2_O_2_, in cultured mammalian cells ^59–62^. To monitor Sima activity in the adult midgut, we reared flies carrying the hypoxia-inducible reporters^63^ *lactate dehydrogenase* (*ldh)-Gal4>UAS-nlsGFP* or *ldh-Gal4>UAS-srcGFP* in control, normoxic conditions and in different damage regimes (Fig. 4a-d; Extended data Fig. 5). These Hif-1*α*/Sima-responsive reporters were expressed at low levels in both midgut epithelial cells and the trachea in baseline conditions. They were strongly induced in both the epithelium and the trachea upon *P.a.* infection or oxidative damage by H_2_O_2_ or PQ (Fig. 4a-d; Extended data Fig. 5). To assess whether *ldh* was induced transcriptionally, we performed RT-qPCR analysis in whole midguts of wild-type *w*^1118^ flies under the same conditions. We found that *ldh* mRNA was induced by *P.a.* infection and even more so by ROS (Fig. 4m). Co-localization experiments with different reporters of midgut cell populations showed that *ldh* transcription was activated in ISCs, EBs and ECs, and rarely in EEs. Importantly, *ldh-Gal4>UAS-srcGFP* highlighted the terminal branches of the intestinal tracheal cells (Fig. 4a-d). Therefore, we surmise that Sima is induced in both the intestinal epithelium and the visceral trachea in response to infection and oxidative stress, suggesting that it may act in both tissues to control tracheal remodeling and intestinal proliferation.

**Figure 4.**
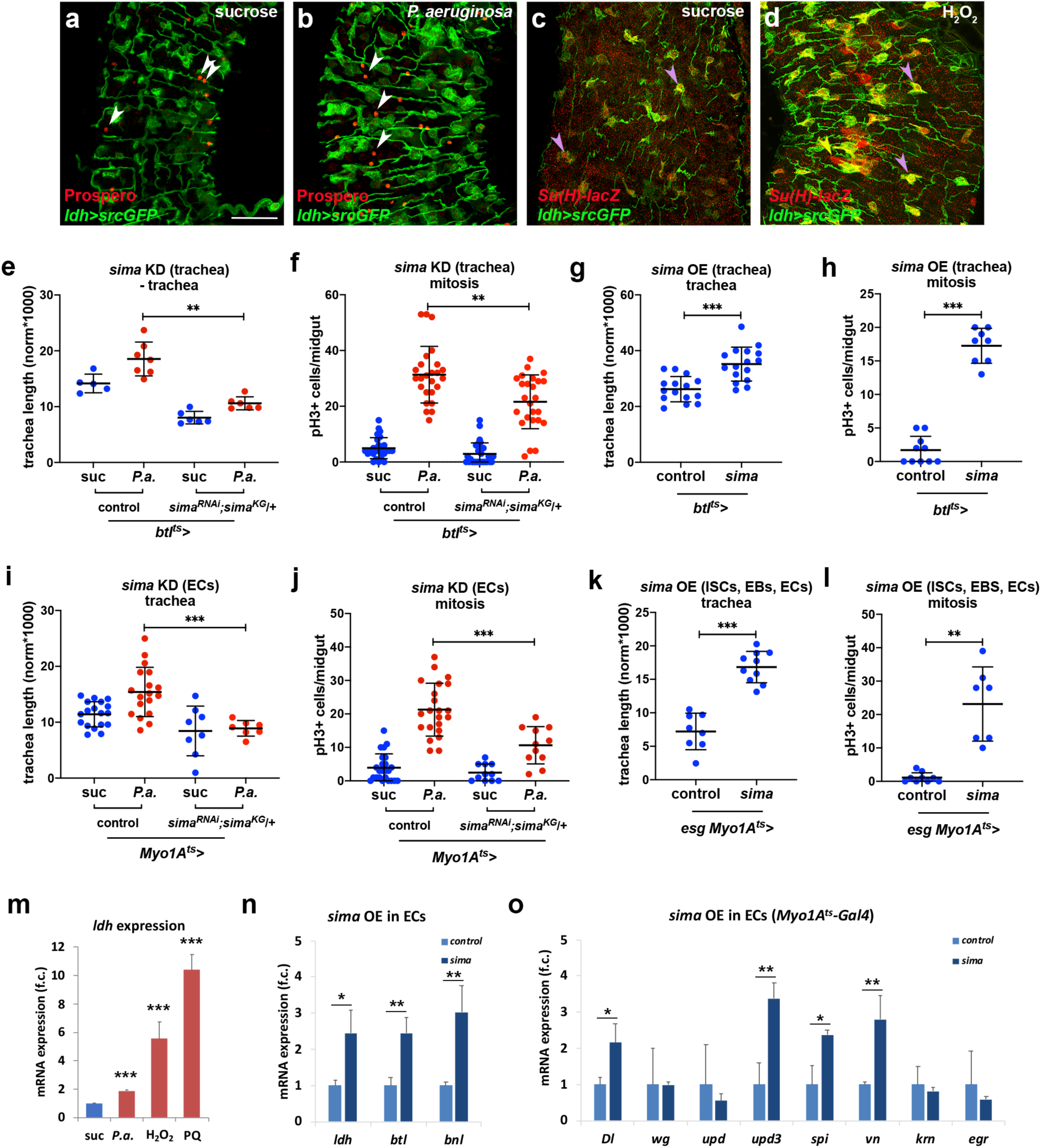
The Hif-1α/Sima is expressed in the midgut epithelium and the visceral trachea, and is necessary and sufficient for ISC proliferation and TTC remodeling. **a-b.** Uninfected (a) and *P.a.*-infected (b) *ldh-Gal4>UAS-srcGFP* midguts (R5 region) with Prospero-labeled (red) EEs. **c-d.** Control (c) and H_2_O_2_-fed (d) midguts (R2 region) with *Su(H)-lacZ*-labeled (red) EBs. Purple arrowheads indicate *ldh*-positive Su(H)-positive EBs and white arrowheads indicate *ldh*-negative Prospero-positive EEs. **e-f.** Quantification of TTC branching (e) and mitosis (f) with or without *P.a.* infection in trachea-specific *sima* knockdown in the background of heterozygous *sima^KG^.* g-h. Quantification of TTC branching (g) and mitosis (h) upon trachea-specific *sima* overexpression in baseline conditions. **i-j.** Quantification of TTC branching (i) and mitosis (j) with or without *P.a.* infection upon EC-specific *sima* knockdown in the background of heterozygous *sima^KG^.* k-l. Quantification of TTC branching (k) and mitosis (l) upon trachea-specific *sima* overexpression in baseline conditions. **m.** RT-qPCR for *ldh* mRNA expression in *w*^1118^ whole adult midguts upon different stresses. **n.** RT-qPCR analysis of *btl*, *bnl* and *ldh* upon EC-specific *sima* overexpression. **o**. RT-qPCR analysis of *Dl, wg, upd, upd3, spi, vn, krn*, and *egr* upon EC-specific *sima* overexpression. The average and standard deviation of biological triplicate experiments are plotted in m-o. Scale bars: 37.5 μm in a-d. Error bars correspond to standard deviation. Statistical significance (t-tested or U-tested, when n<10): ns, not significant, * 0.01<*p*≤0.05, ** 0.001<*p*≤0.01 and *** *p*≤0.001.

To assess whether Hif-1α/Sima induction is necessary for gut tracheal remodeling and ISC mitoses, we performed tissue-specific loss- and gain-of-function tests with *sima*. To achieve an effective *sima* knockdown, we inactivated *sima* by tissue-specific RNAi in a heterozygous *sima^KG^* mutant background ^64^, either in the tracheal cells or in ECs. Upon *P.a.* infection, both tracheal coverage and ISC proliferation rates were reduced by *sima* depletion (Fig. 4e-f,i-j; Extended data Fig. 6). To assess if *sima* is sufficient for tracheal remodeling and intestinal regeneration in baseline conditions, we overexpressed wild-type *sima* ^11^ specifically in adult tracheal cells, as well as in progenitors and ECs, using *btl^ts^-Gal4* and *esg^ts^-Gal4 Myo1A^ts^-Gal4*, respectively. We found that overexpressed *sima* was sufficient to induce both tracheal remodeling and ISC proliferation in the absence of infection (Fig. 4g-h,k-l). Interestingly, *sima* gain of function was also associated with increased expression of the *bnl* and *btl* transcripts, suggesting that *bnl* and *btl* expression is downstream of *sima* activation in the midgut (Fig. 4n). Furthermore, EC-specific *sima* overexpression led to increased expression of known ISC mitosis regulators, such as genes encoding the Notch ligand Delta (Dl), the Jak/Stat ligand Unpaired 3 (Upd3), and the EGFR pathway ligands Spitz (Spi) and Vein (Vn) (Fig. 4o). Thus, Hif-1α/Sima has a key role in midgut TTC remodeling and damage repair acting both in the epithelium and the intestinal trachea.

### Increased tracheation induces ISC proliferation in normoxia, but not in hypoxia

In addition to bacteria and ROS, low oxygen tension (hypoxia) is well known to activate the Hif-1α, and to effect remodeling of oxygen-carrying blood vessels and trachea, in mammals and insects, respectively ^65, 66^. To assess the effect of oxygen tension on intestinal tracheogenesis in adult *Drosophila*, we subjected wild-type flies to hypoxia (5% O_2_) for 12 hrs and compared them with flies reared in normoxia (21% O_2_). We found that intestinal tracheal coverage, but not the number of TTCs, was significantly increased in flies subjected to hypoxia (Fig. 5a-c, g). If hypoxia induces tracheal remodeling and tracheal remodeling, in turn, promotes ISC mitoses, we might expect increased mitoses in hypoxia-treated flies. In contrast, we found that upon hypoxia, intestinal mitosis was significantly reduced in wild-type flies (Fig. 5d). Reduced mitosis in hypoxia was not the result of midgut progenitor death, because *esg+* progenitor apoptosis was not observed in hypoxic intestines (Fig. 5e-f) and the number of *esg+* progenitors was unchanged between treatments (Fig. 5h). Hypoxia-induced tracheal expansion was also associated with induction of reporters for *bnl* (*bnl-Gal4>srcGFP*) and *sima* (*ldh-Gal4>srcGFP*) in the epithelial cells, and in the epithelium plus the intestinal trachea, respectively (Fig. 5i-l). In addition, flies subjected to hypoxia for 4 hrs exhibited increased mRNA expression of *bnl* and *ldh* in their midguts. Although *btl* was not induced upon a 4 hr hypoxia treatment, 12 hrs in hypoxia-induced *btl* in the adult midgut by approximately 2-fold (Fig. 5q). Therefore, although hypoxia induces intestinal tracheal remodeling, it does not promote increased ISC mitosis, uncoupling the two processes.

**Figure 5.**
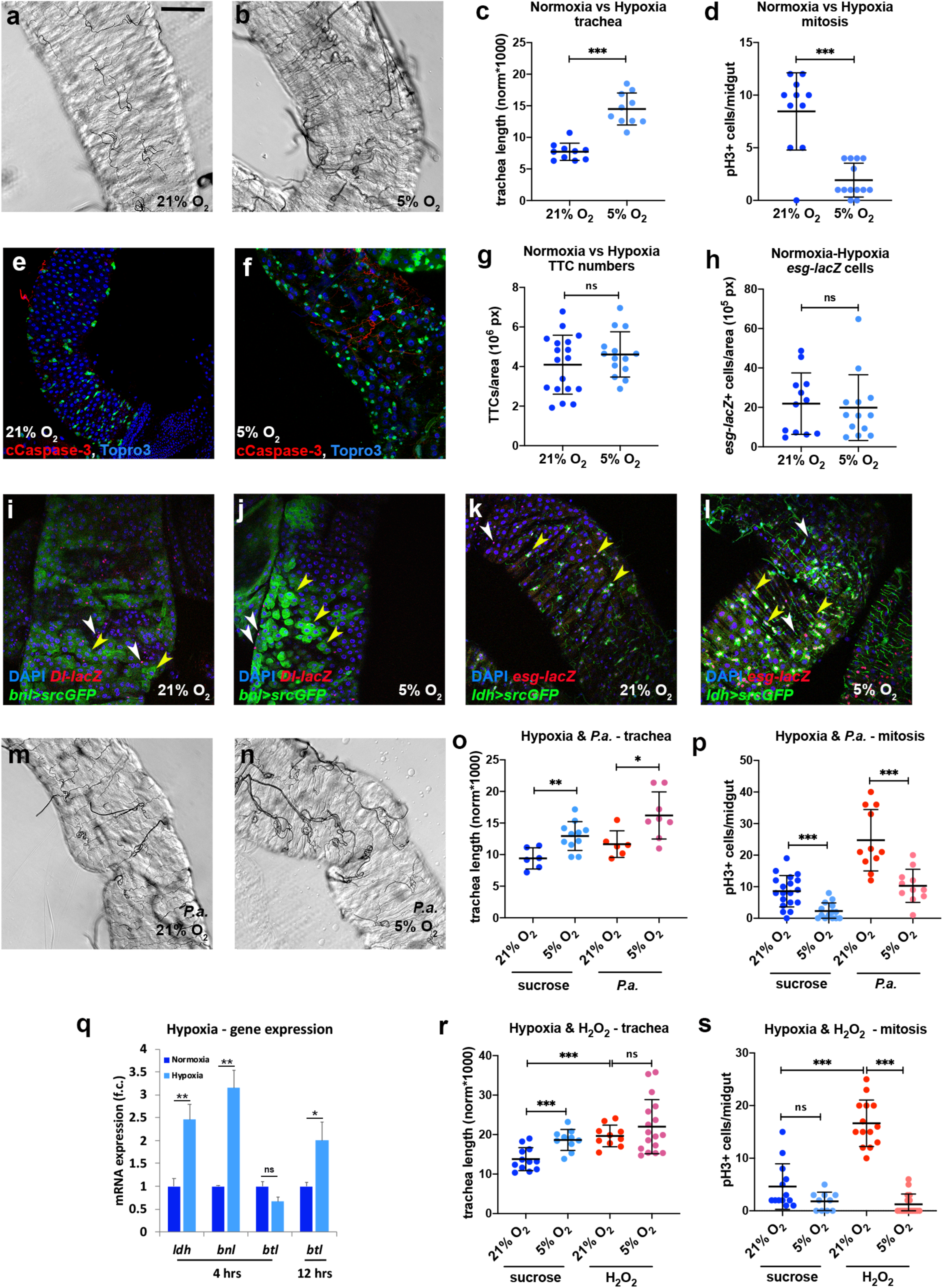
Hypoxia induces TTC branching, but not ISC proliferation. **a-b.** Brightfield images of the midgut TTCs (R5 region) of wild-type (*w*^1118^) uninfected flies reared in normoxia (21% O_2_) vs. hypoxia (5% O_2_). **c-d.** Quantification of TTC branching (c) and mitosis (d) in normoxia vs. hypoxia. **e-f.** *esg-Gal4>UAS-srcGFP* (ISCs+EBs, green) midguts (R5 region) of flies reared in normoxia (e) and hypoxia (f) stained for cleaved Caspase-3 (red) and Topro3 (blue) staining all nuclei. **g-h**. Quantification of TTCs (g) and *esg-lacZ+* progenitors (h) in normoxia and hypoxia. **i-j.** Normoxia-(i) and hypoxia-reared (j) *ldh-Gal4>UAS-srcGFP* (Sima reporter, green) midguts with *esg-lacZ* (red) labeling the ISCs and EBs and DAPI (blue) staining all nuclei. Yellow arrowheads indicate *bnl*-positive and *esg*-positive progenitors and white arrowheads indicate *bnl*-negative and *esg*-positive progenitors. **k-l**. Normoxia-(k) and hypoxia-reared (l) *ldh-Gal4>UAS-srcGFP* (Sima reporter, green) midguts with *esg-lacZ* (red) labeling the ISCs and EBs and DAPI (blue) staining all nuclei. Yellow arrowheads indicate *bnl*-positive and *esg*-positive progenitors and white arrowheads indicate *bnl*-negative and *esg*-positive progenitors. **m-n.** Brightfield images of the midgut TTCs (R5 region) of wild-type (*w*^1118^) *P.a.-*infected flies reared in normoxia (21% O_2_) vs. hypoxia (5% O_2_). **o-p.** Quantification of TTC branching (o) and mitosis (p) of uninfected and *P.a.*-infected flies in normoxia and hypoxia. **q..** RT-qPCR analysis of *btl*, *bnl* and *ldh* in normoxia vs. hypoxia. The average and standard deviation of biological triplicate experiments are shown. **r-s.** Quantification of TTC branching (r) and mitosis (s) of untreated vs. H_2_O_2_-fed flies in normoxia and hypoxia. Scale bars: 75 μm. Error bars correspond to standard deviation. Statistical significance (t-tested or U-tested, when n<10): ns, not significant, * 0.01<*p*≤0.05, ** 0.001<*p*≤0.01 and *** *p*≤0.001.

To assess the relationship between infection and hypoxia, both of which induce tracheal remodeling but affect ISC mitoses oppositely, we orally infected flies with *P.a.* while subjecting them to hypoxia. Although tracheal branching was increased in treated flies (Fig. 5m-o), hypoxia significantly suppressed the mitotic response to *P.a.* infection (Fig. 5p). To rule out the possibility that hypoxia affected the damaging capacity of the *P.a.* bacteria, we also tested tracheal branching and mitosis of H_2_O_2_-treated flies upon hypoxia. In this case, regardless of the increased tracheal coverage, mitoses were also suppressed by hypoxia (Fig. 5r-s). These results indicate that although increased trachea is able to induce ISC proliferation, the trachea does so only in the presence of oxygen (*i.e.* in normoxic conditions). This implicates oxygen availability as a necessary component for tracheal induction of ISC mitoses.

### Bacterial ROS-generators induce ISC proliferation via the tracheal niche

Enteric *Pseudomonas* infection phenocopies oxidative stress with regards to intestinal regeneration and tracheal remodeling. In addition, *Pseudomonas*, as extracellular pathogens, damage tissues via secreted virulence factors. The *P.a.* quorum-sensing MvfR pathway is a master regulator of virulence. Genes involved in phenazine production, including *phzS* and *phzM*, control pyocyanin, a ROS-generating secreted virulence factor that provides the blue hue in the *P.a.* cultures ^67, 68^. To assess the role of *P.a.* pyocyanin in *Drosophila* intestinal regeneration and tracheogenesis, we fed adult wild-type flies with bacterial culture supernatants from wild-type *P.a.* (strain PA14) and MvfR, PhzM and PhzS mutants (isogenic mutants in the PA14 background ^69^). All *P.a.* mutants tested led to significant reductions of tracheal remodeling (Fig. 6a-g) compared to the wild type PA14 control, and this was accompanied by significant reductions in ISC mitoses (Fig. 6h). Since pathogenic bacteria are known to induce intracellular ROS via the Dual Oxidase (Duox) ^32, 70^, we assessed the requirement for *Duox* in wild type *P.a.-*induced intestinal tracheal remodeling and ISC mitosis. To assess if epithelial or tracheal *Duox* is involved in the infection response, we knocked down *Duox* in the epithelial ECs or the trachea via *Myo1A^ts^-Gal4* and *btl^ts^-Gal4*, respectively. We found that *Duox* was necessary in both the epithelium and the trachea for induction of tracheal remodeling and mitosis (Fig. 6i-l). Thus, bacterial ROS-generating agents present in the *P.a.* supernatant are necessary for tracheal remodeling and intestinal mitosis and act on both the epithelium and the intestinal trachea.

**Figure 6.**
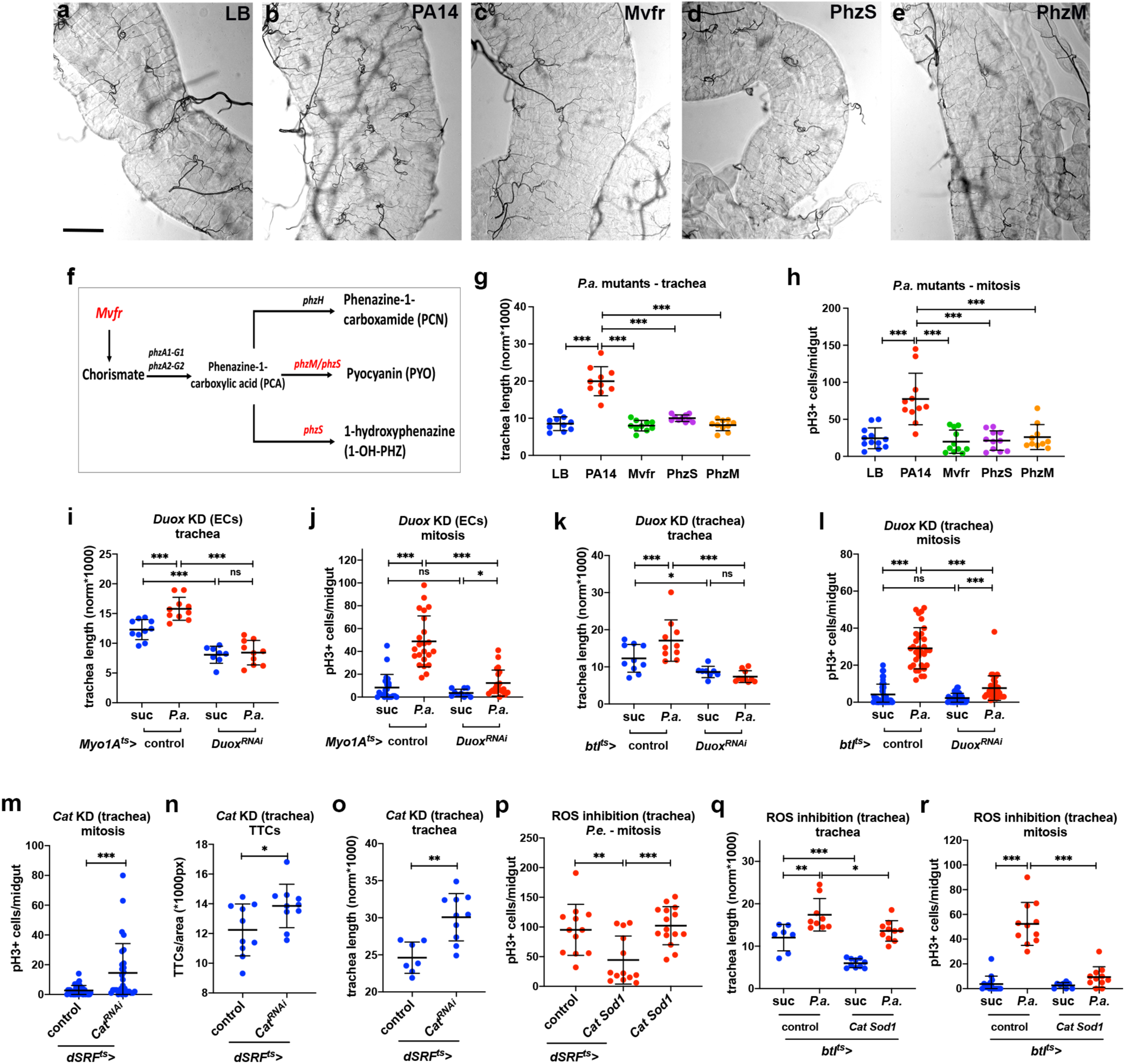
Stress-responsive increased TTC remodeling is ROS dependent. **a-e.** Brightfield images of the midgut TTCs (R5 region) of wild-type (*w*^1118^) flies fed on LB (a, negative control) and *P.a.* culture supernatants of the virulent wild-type PA14 strain (b, positive control), the quorum-sensing mutant MvfR (c), and the phenazine mutants PhzS (d) and PhzM (e). **f.** Schematic of the *P.a.* quorum-sensing and phenazine production pathway. **g-h.** Quantification of TTC branching (g) and midgut mitosis (h) upon *P.a*. culture supernatant feeding. **i-j.** Quantification of TTC branching (i) and midgut mitosis (j) upon EC-specific *Duox* knockdown with or without *P.a*. infection. **k-l**. Quantification of TTC branching (k) and midgut mitosis (l) upon trachea-specific (via *btl^ts^-Gal4*) *Duox* knockdown with or without *P.a*. infection. **m-o**. Quantification of midgut mitosis (m), TTC number (n) and TTC branching (o) upon trachea-specific (via *dSRF^ts^-*Gal4) *Cat* knockdown in uninfected flies. **p**. Quantification of midgut mitosis upon trachea-specific (via *dSRF^ts^-*Gal4) *Cat* and *Sod1* overexpression in *P.e.-*infected flies. **q-r.** Quantification of TTC branching (q) and midgut mitosis (r) upon trachea-specific (via *btl^ts^-Gal4*) *Cat* and *Sod1* overexpression with or without *P.a*. infection. Scale bars: 75 μm. Error bars correspond to standard deviation. Statistical significance (t-tested): ns, not significant, * 0.01<*p*≤0.05, ** 0.001<*p*≤0.01 and *** *p*≤0.001.

It is known that ROS induce ISC mitoses and cell turnover in the fly’s intestinal epithelium ^46, 71^. However, no report thus far has indicated any direct effects of ROS on the intestinal trachea. Nevertheless, we find that bacterial secreted ROS-generators and chemical ROS affect the intestinal trachea (Fig. 1a-l, Fig. 6a-h and Extended data Fig. 1) via the Hif-1α/Sima transcription factor. To address the role of tracheal ROS in the process of damage-induced tracheal remodeling and mitosis, we genetically modulated ROS levels or activity directly in the trachea and measured both tracheation and ISC proliferation rates. Trachea-specific RNAi against the antioxidant *Catalase A* (*Cat*) in baseline uninfected conditions showed a three-fold higher ISC mitotic index as compared to controls. This was accompanied by increased visceral tracheole branching, but no differences in the number of TTCs (Fig. 6m-o). This suggests that ROS indeed exist in tracheal branches and affect the epithelium. On the contrary, when antioxidants, *Cat* and *Superoxide dismutase 1* (*Sod1*), were overexpressed in the adult trachea, the infection-induced ISC proliferation was severely restricted (Fig. 6p,r). Trachea-specific overexpression of *Cat* and *Sod1* also led to a significant reduction in tracheal branching (Fig. 6q). Thus, our results suggest that flies remodel their intestinal tracheal branch network in response to ROS-mediated tissue damage that induces Hif-1α/Sima in both the midgut epithelium and the trachea. In normoxia, increased tracheal branches increase oxygen supply, which in turn is necessary for the efficient regenerative response of ISCs.

### Stem cell-derived tumors induce and require tracheal arborization

Neoangiogenesis is a hallmark of mammalian cancer ^72^. While differentiation of tumor cell into trachea has been alluded to ^54, 73^ and tumor-trachea association has been reported ^30^ previously, no study thus far has examined the role of the *Drosophila* trachea in tumorigenesis. To investigate whether *Drosophila* ISC-derived tumor development is associated with increased tracheal branching (“neo-tracheogenesis”), we generated MARCM clones expressing the *Ras^V12^* oncogene in the adult midgut. We observed increased density of tracheoles in the tumor compared to the surrounding tissue (Fig. 7a). Tracheal length measurements of intestines carrying control wild type versus *Ras^V12^* MARCM clones indicated a clear increase in tracheation in the *Ras^V12^* (Fig. 7b). To test the requirements of the trachea for the tumor growth, we transplanted progenitor derived *Ras^V12^ Notch^RNAi^* tumor masses labeled with GFP or RFP into recipient flies with mCherry- or GFP-labeled trachea, respectively, and found that the wild-type trachea of the recipient was recruited to the transplanted tumor (Fig. 7c-d). In addition, since Btl is necessary for tracheal growth, we transplanted *Ras^V12^ Notch^RNAi^* tumorous cell masses into recipient wild-type flies or flies with RNAi-mediated knockdown of *btl* in the trachea. We found that 85% of the transplanted tumors could survive and grow when transplanted in wild-type recipients. However, only 45% could survive in the trachea-specific *btl^RNAi^-*expressing recipients, and the tumor masses were substantially smaller (Fig. 7e). In addition, we generated transgenic flies bearing stem cell-derived intestinal tumors. The tumors were generated by expressing either a dominant negative form of the tumor suppressor Notch (*Notch^DN^*), an activated form of the Ras GTPase (*Ras^V12^* or *Ras^Q13^*) (Fig. 7; Extended data Fig. 7-8) in adult intestinal progenitor cells. In all genotypes we observed increased tracheal branching either by fluorescent tracheal labeling (Fig. 7f-g, k-l, Extended data Fig. 7-8) or by monitoring the tracheal tubes in midguts by brightfield microscopy (Fig. 7a-b, Fig. 8a). To achieve simultaneous labeling of both the tumors and the intestinal trachea, we combined the Gal4/Gal80^ts^/UAS system^50, 74^ with the QF/QUAS system^52^, so that the tumors were visualized with eGFP and the trachea with a membrane-targeted mtd-Tomato. Then, we performed time-course analysis for the *Notch^DN^* and the *Ras^V12^* tumors to simultaneously visualize their development and their tracheal branching. We found that *Notch^DN^* caused relatively slow, regional tumorigenesis, so that tumors developed gradually and were clearly visible in the R4a (P1) region of the midgut^75, 76^ after 4 days of induction (Extended data Fig. 7). The *Ras^V12^*-expressing tumors, on the other hand, developed rapidly. Their growth was noticeable after 1 day of induction (Fig. 7g), and eGFP-positive cells almost covered the midgut 3 days after tumor initiation (Extended data Fig. 8). In both tumor genotypes, tumor growth correlated with an expansion of TTC branching. Quantification analysis of the intestinal trachea (tracheal branch length normalized to the midgut surface in the R4a region for *Notch^DN^* and throughout the midgut for *Ras^V12^*) showed significant and gradual increases in the tumorous midguts (Fig. 7f-h, Extended data Fig. 7a-d; e-g; Extended data Fig. 8). In addition, the tumor-bearing midguts encompassed more mitotic cells than those of the control cohorts (Fig. 7i, Extended data Fig. 7g). Similarly, double *Ras^V12^ Notch^RNAi^* tumors exhibited increased tracheal branching (measured as the number of tertiary tracheal branches in R5), as well as mitoses (Fig. 8a,c-d). These results, therefore, suggest that the visceral trachea is induced by the tumor and is necessary for its growth, and that tracheal branches are needed for this process.

**Figure 7.**
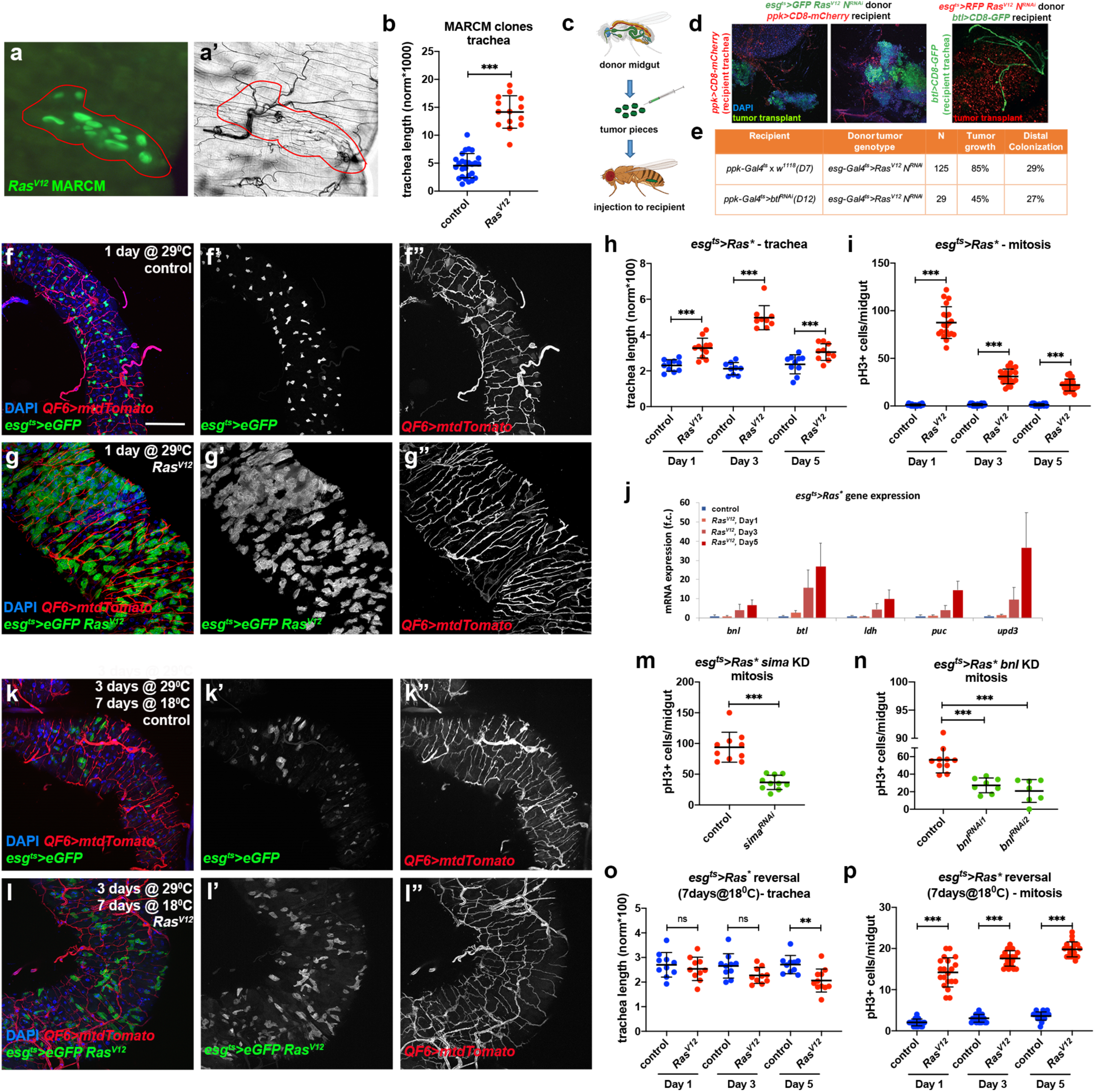
Midgut ISC-derived tumors induce TTC branching required for tumor growth. **a**. Midgut *Ras^V12^* MARCM clone (green) and corresponding brightfield image (a’) of the trachea. **b**. Quantification of TTC branching in tumorous *Ras^V12^* clone-bearing midguts vs control clone-bearing midguts. **c**. Schematic of the transplant experimental process. **d.** Fluorescently-labeled midgut tumor pieces from a donor fly are transplanted to a recipient fly with fluorescently-labeled trachea. Images showing GFP+ green tumors (first two images) recruiting CD8-mCherry marked tracheae from the recipient. The third image shows RFP+ (red) tumor mass overlayed with CD8-GFP+ (green) recipient trachea. **e**. Table showing comparison of the growth ability and survival of transplanted tumors in wild-type or trachea-defective (*ppk-Gal4>UAS-btl^RNAi^*) recipients. **f-g**. The R5 region of control *esg-Gal4 UAS-eGFP tub-Gal80^ts^* (f) and *esg-Gal4 UAS-eGFP tub-Gal80^ts^*>*UAS-Ras^V12^*-tumor (g) bearing midguts with concomitant expression of *QF6>QUAS-mtdTomato* (red) to label the trachea reared for 1 day at 29^°^C. DAPI (blue) was used to label all midgut nuclei. f’-g’, and f’’-g’’ show the individual channels for the eGFP and the Tomato-labeled trachea, respectively. **h-i**. Quantification of TTC branching (h) and midgut mitosis (i) in control and *Ras^V12^*-tumor bearing flies during a time-course analysis at 1, 3, and 5 days post-tumor induction. **j**. qRT-PCR analysis of *btl*, *bnl*, *ldh, puc and upd3* upon *Ras^V12^* tumor progression. **k-l.** Control genotype and *Ras^V12^* tumor reversal (3 days at 29^°^C followed by 7 days at 18^°^C). k’-k” and l’-ll’’ show the individual channels for eGFP and the Tomato-labeled trachea, respectively. **m-n**. *Ras^Q13^* tumor mitosis in the presence or knockdown of *sima* (m), and *bnl* (n). **o-p**. Quantification of TTC branching (o) and midgut mitosis (p) in control and *Ras^V12^* following reversal of the tumors after 1, 3, and 5 days post tumor initiation. Error bars correspond to standard deviation. Scale bars: 75 μm in f-g, k-l. Statistical significance (t-tested): ns, not significant, * 0.01<*p*≤0.05, ** 0.001<*p*≤0.01 and *** *p*≤0.001.

**Figure 8.**
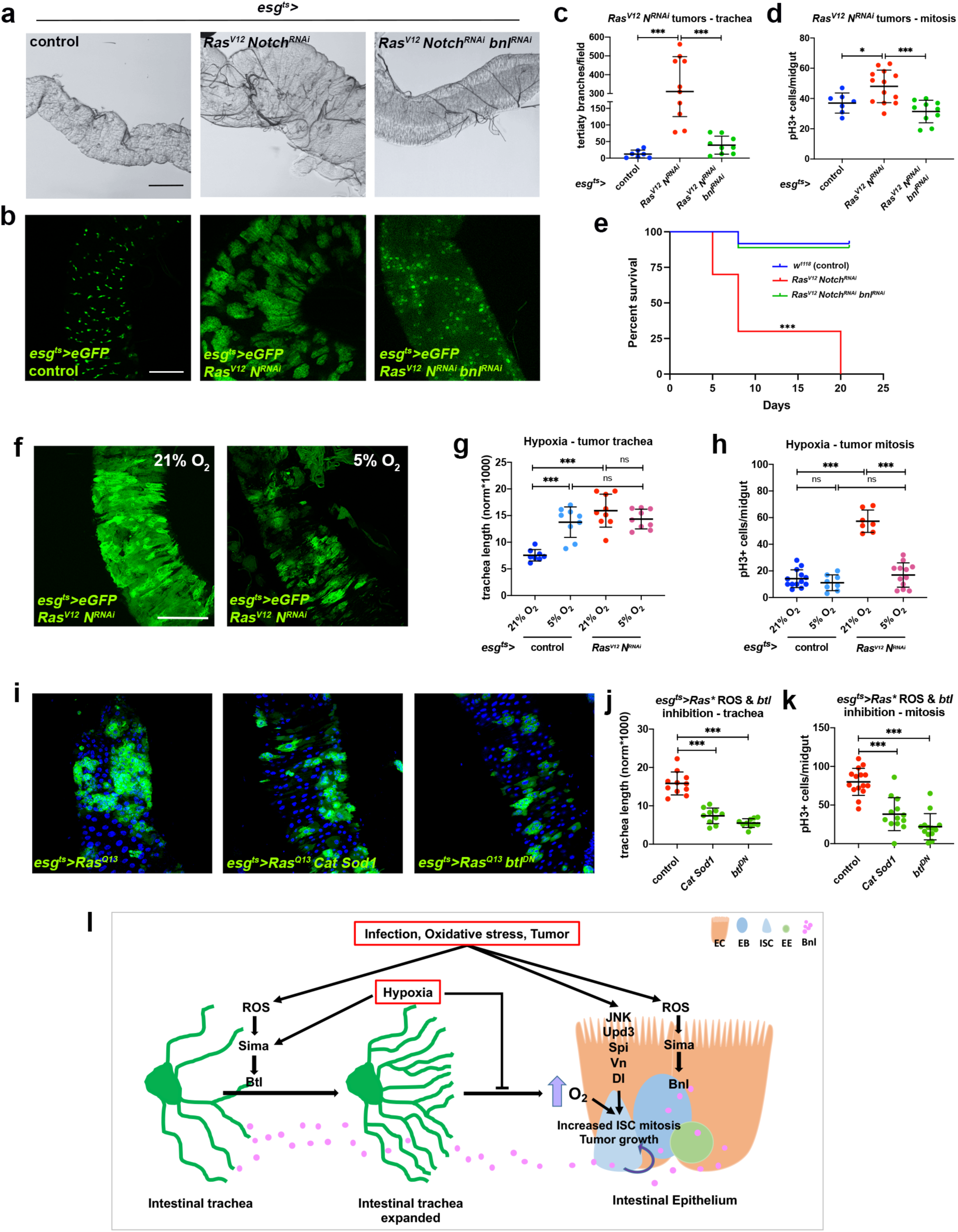
Tumor-induced increased tracheal coverage, oxygen supply, and ROS are necessary for tumor growth. **a-d**. Brightfield images of the TTCs (a) and fluorescent images of the GFP^+ve^ (b) progenitor or progenitor derived tumor cells in control midguts, midguts bearing *esg-Gal4 UAS-eGFP tub-Gal80^ts^*>*UAS-Ras^V12^ UAS-Notch^RNAi^* tumors, and *esg-Gal4 UAS-eGFP tub-Gal80^ts^*>*UAS-Ras^V12^ UAS-Notch^RNAi^* co-expressing *UAS-bnl^RNAi3^*. Quantification of TTC branching (c) and mitosis (d) in control, *Ras^V12^ Notch^RNAi^* and *Ras^V12^ Notch^RNAi^ bnl^RNAi^* tumor-bearing flies. **e.** Percent survival of control, *Ras^V12^ Notch^RNAi^* and *Ras^V12^ Notch^RNAi^ bnl^RNAi^* tumor-bearing flies. **f-h**. Images showing growth of *Ras^V12^ Notch^RNAi^* tumors (f), quantification of TTC branching (g) and midgut mitosis (h) in control and *Ras^V12^ Notch^RNAi^* tumor-bearing flies in normoxia vs. hypoxia. **i-k**. Images showing tumorous midguts (R5 region) with progenitor derived *Ras^Q13^* tumors (green) with *Cat* and *Sod1* overexpression or *btl* knockdown. Quantification of TTC branching (j) and midgut mitosis (k) in the same genotypes. Scale bars: 200 μm in a, 100 μm in b and 75 μm in f and i. **m.** Model of the intestinal trachea-midgut communication during damage-induced regeneration. Error bars correspond to standard deviation. The Kaplan-Meier method was used to test significance in e. For all others, statistical significance (t-tested): ns, not significant, * 0.01<*p*≤0.05, ** 0.001<*p*≤0.01 and *** *p*≤0.001.

To assess whether tumors induce *btl*, *bnl* and *sima* similar to infection and oxidative damage, we performed RT-qPCR analysis of tumorous midguts and compared them to wild-type controls. We found that *Ras^V12^* tumors upregulate, as expected, damage-induced genes, such as *puckered* (*puc*) and *upd3*, and that *btl*, *bnl* and *ldh* (the reporter gene for Sima activity) were also upregulated (Fig. 7j). To assess the requirement for *sima* induction in tumors, we knocked down *sima* via RNAi in the tumor cells. We found that co-expression of *sima^RNAi^* led to reduced number of tumor mitoses (Fig. 7m). To assess the requirement for *bnl*, we knocked down *bnl* in the tumor cells using RNAi, wherein tumor size, mitoses, as well as tracheal coverage were found reduced (Fig. 7n and Fig. 8a-d). These results suggest that *bnl* is also required for tumor growth, mitoses and increased tracheal branching around the tumors (Fig. 7n and Fig. 8a-d). Further, we performed survival experiments on tumor-bearing flies with or without *bnl^RNAi^* expression in the tumors. We found that the survival of the tumor-bearing flies was significantly increased, when *bnl^RNAi^* was expressed in the tumors (Fig. 8e). Collectively, these results suggest that the activation of *sima* and *bnl* in intestinal tumors induces tracheal outgrowth, which enhances tumor growth.

To assess whether induction of tracheal remodeling is a reversible feature of growing tumors, we discontinued *Ras^V12^* expression 1, 3 and 5 days after induction by shifting the flies from 29^°^C to 18^°^C (Fig. 7k-l). We found that *Ras^V12^* tumor growth and associated tracheal growth could be effectively reversed by switching *Ras^V12^* expression off for 7 days, 1, 3, or 5 days post oncogene induction (Fig. 7o-p; also compare to Fig. 7h-i). Further, since increased tracheal branching was associated with increased ISC proliferation only in normoxia, we hypothesized that tumor growth would be reduced in hypoxia. To test this, we reared tumor-bearing flies under hypoxia. Indeed, both tumor size and ISC proliferation were reduced by hypoxia, although no significant difference in intestinal tracheal coverage was observed between normoxia and hypoxia (Fig. 8f-h). We surmise that, during tumorigenesis, the visceral trachea expands TBs to accommodate the extra metabolic needs of the growing tumors and to sustain tumor mitoses. In hypoxia, however, due to the limited oxygen availability to the tumor, regardless of expanded tracheation, tumor growth is reduced. Furthermore, to assess the involvement of ROS in tumor growth and tracheal remodeling, we stained *Ras^V12^* tumorous midguts with dihydroxyethidium (DHE) and confirmed previous observations ^46^ for the presence of tumor-induced ROS (Extended Data Fig. 9). To test the tumor-autonomous role of ROS in intestinal tracheal branching, we inhibited ROS in the tumor by overexpressing *Cat* and *Sod1*. We found that tumor growth, mitosis and tracheole branching were reduced (Fig. 8i-k). Finally, to test if *btl* has a non-tracheal role in the growing tumors, as previously suggested ^73^, we inactivated *btl* by overexpressing *btl^DN^* in the *Ras^V12^* tumorous midguts. We found that *btl* is necessary for tumor growth, mitosis and tracheole branching (Fig. 8i-k). Thus, ROS and *btl* are necessary for tumor mitosis and tracheole remodeling.

### Conclusions - Discussion

Angiogenesis is the process by which oxygen-carrying blood vessels are remodeled physiologically during development, wound healing or pregnancy ^77^. This process can go awry during inflammatory diseases and is recognized as one of the hallmarks of cancer, since tumor cells induce neoangiogenesis to achieve a growth advantage and become metastatic ^72, 78^. A plethora of studies in cell or tumor culture, as well as vertebrate models have established the link between tumor growth and angiogenesis ^79–81^. In this context, hypoxia is an important inducer of angiogenesis, working via the Hif-1α pathway ^82, 83^. Despite the fact that several tumor angiogenesis regulators have been identified, including VEGF, PDGF, and FGF ^80, 84^, we know little about the specific environmental factors that contribute to their induction and whether they are also induced in cases other than tumorigenesis, for example, by infection or oxidative stress.

We used the adult *Drosophila* midgut as a model to study the signals involved in tissue communication between the intestinal epithelial cells and the visceral trachea in conditions of homeostasis, midgut damage, as well as tumorigenesis. We found that the adult intestinal terminal tracheal cells (TTCs) are dynamic. They respond to ROS-generating secreted bacterial factors, ROS-inducing chemicals and tumors by extensively expanding their terminal branches (TBs) and overall tracheal coverage of the midgut. This damage-induced TB expansion phenocopies hypoxia-induced TB expansion in the adult midgut (Figures 1 and 5), suggesting a role for increased tracheal coverage to maximize oxygen supply both in response to oxidative damage and hypoxic conditions. The plasticity of the adult trachea in response to insulin/oxygen-sensitive neurons in the *Drosophila* alimentary canal has been shown earlier ^29^. In this study, intestinal tracheal arborization mostly of the hindgut was found to depend on insulin/oxygen sensitive neurons that transport both local and systemic insulin- and VIP-like neuropeptides. This neuron-controlled tracheal branching can affect metabolic changes such as lipid storage. Whether the neuronal network also regulates the damage-induced tracheal branching and ISC proliferation is an interesting and open question, that should be pursued in further studies.

Branching morphogenesis is triggered by cell-cell communication that involves tracheal cells expressing the FGFR/Btl on their surface following guidance cues from neighboring cells expressing the FGF/Bnl during development ^1^. Not much is known, however, about the adult tracheal remodeling, and the role of non-tracheal functions of Btl during tracheal remodeling. A study focusing on the fly wing disc has shown that hypoxic tumor cells stabilize Sima and induce *bnl.* A subset of these tumor cells differentiates into tracheal cells expressing Trachealess (Trh), which is a known transcriptional inducer of *btl* ^85^, in ways reminiscent of pathological mammalian angiogenesis ^54^. In addition, EB-derived midgut tumors caused by *Sox21a* deletion have been shown to induce *btl* unconventionally in the tumor ^73^. A recent study on ISC-derived *Apc-Ras* tumors has shown that tumor cells expressing Snail undergo epithelial-mesenchymal transition, and during their migration through the basal lamina, they envelope the intestinal trachea using it as a scaffold. Interestingly, upon 10 rounds of transplants, these tumors lose their mesenchymal character and associate with tracheal cells. The authors suggested a role of the trachea for tumor oxygenation to support its growth, but they did not test the tracheal requirement directly ^30^. Our study confirms that the trachea actively regulates tumor growth and is an essential part of niche modification by the tumor for its growth and survival.

We found that *btl* is expressed and induced in the visceral tracheal cells, and its tracheal expression is necessary for tracheal maintenance, remodeling, and ISC mitosis. Similarly, the ligand for Btl, Bnl is found in the intestinal epithelial cells, where the *bnl* gene is induced upon damage. Epithelial *bnl* is necessary for tracheal remodeling and ISC mitosis. Tracheal *btl* and epithelial *bnl* overexpression were also sufficient to promote both tracheal remodeling and regeneration. These data are in agreement with the expected function of the Btl/Bnl pathway in tracheal remodeling. Surprisingly, although we could not detect *btl* in the epithelium or *bnl* in the trachea by reporter expression, we found that epithelial *btl* and tracheal *bnl* inhibition led to impaired tracheal remodeling and ISC mitosis in response to damage (Fig. 2; Fig. 3). This indicates that epithelial *btl* and tracheal *bnl* are necessary for the damage-induced regeneration response which underscores the Btl receptor’s novel role in intestinal mitosis. It is worth noting that we observed an inconsistency in the tracheal *bnl* knockdown experiments: when *bnl^RNAi^* was driven by *btl^ts^-Gal4*, it affected both mitosis and tracheal remodeling, but when it was driven by *dSRF^ts^-Gal4*, it led to impaired mitosis, but no significant effect on tracheal coverage was observed. These results could be explained by the different strength of the two drivers used. The *btl-Gal4* is expressed broadly and strongly in all the tracheal cells, whereas the *dSRF-Gal4* is considered a TTC-specific driver and is weaker than the *btl-Gal4*. Strikingly, overexpression of *btl* in the epithelium and *bnl* in the trachea were not sufficient to induce tracheole remodeling or ISC mitosis (Fig. 2; Fig. 3). This is in contrast to the overexpression effects in the tissues where their expression is easily detectable (i.e *btl* in the trachea and *bnl* in the epithelium). Since there is no evidence supporting a cis interaction between Bnl and Btl in the tracheal cells or elsewhere, we assume that the interaction between the ligand and the receptor involves different cells. Our loss of function results indicate that *btl* and *bnl* are required in cells that express them strongly (the trachea and epithelium), as well as other midgut cells. On the contrary, our gain of function experiments show that *btl* is only sufficient in the trachea and *bnl* is only sufficient in the epithelium to induce the regenerative response. These results suggest that the function of Btl and Bnl in different intestinal cells contributes to the plasticity of the tissue and its quick response to damage. Tracheal Btl is abundant, and its sensitivity to epithelial Bnl levels, which are variable and arise from multiple sources, directly controls tracheal branching, and as a consequence, results in ISC mitosis. On the other hand, epithelial Btl and tracheal Bnl are present at lower levels in the intestine to fine-tune ISC mitosis and, in turn, indirectly impinge on tracheal remodeling.

Our data uncover a novel role for Hif-1α/FGF signaling in the dynamic, inter-organ control of *Drosophila* intestinal stem cells. Upon midgut damage, both the FGF receptor (Btl) and the FGF ligand (Bnl) are regulated by the Hif-1α transcription factor homolog, Sima. This is reminiscent of the effects of Sima in larvae developing under hypoxia ^11^. In our study, however, the trigger for Sima is ROS in normoxia. ROS are a major byproduct of oxygen metabolism within cells and have an emerging role in regeneration and injury response in both *Drosophila* and mammals ^45, 46, 86, 87^. Interestingly, in mammals ROS regulate angiogenesis in a tissue-specific manner. For example, ROS suppress angiogenesis in diabetic mice but enhance placental angiogenesis during pregnancy ^88, 89^. Epithelial tumors induce ROS-mediated signaling to promote their proliferation in mammals and *Drosophila*^46, 90, 91^. Since we found that bacteria and oxidative agents induced increased tracheal branching, we assessed the contribution of ROS to the midgut tracheal phenotype. Our data indicate that bacterially-secreted ROS-generators or chemical ROS reach the trachea directly to promote TTC remodeling and ensure ISC proliferation.

It has been shown that pathogenic bacteria and commensals induce endogenous ROS-producing NADPH oxidases in the intestinal epithelium, the Dual oxidase (Duox) and the NADPH oxidase (Nox), respectively, as an innate immune mechanism ^92–94^. Our data show that ROS impinge on the remodeling of the intestinal trachea directly, as well as affect regeneration (Fig. 5). It has been shown that upon *P.e.* infection intracellular ROS produced in ECs by Nox and potentially Duox are necessary for intestinal regeneration ^46^. *Duox* or *Nox* mRNAs were not induced upon *P.a.* infection for 48 hrs (not shown), but we cannot exclude the possibility that they are induced at an earlier time-point. When we assessed if H_2_O_2_ and PQ had any effect on *Duox* mRNA, we found that it was significantly reduced, indicating a negative feedback regulation of intracellular ROS production by the excess ingested ROS (not shown). When we tested the requirement for *Duox* in *P.a*.-infected midguts, we found that it was necessary for mitosis and tracheal remodeling in both the epithelial cells and the intestinal trachea (Figure 6i-l). Although *Duox* is highly expressed in barrier epithelia including the larval *Drosophila* trachea ^95^, its tracheal role in regeneration has never been demonstrated. Here, we show for the first time that *Duox* is involved in tracheal remodeling. Taken together, our data show that exogenous ROS and intestinal ROS act coordinately in the trachea and the epithelial cells for regeneration.

As we expected, hypoxia activated the Hif-1α/FGF cascade in the gut and associated trachea, promoting TTC remodeling. However, regardless of the extensive tracheal coverage, hypoxia suppressed ISC mitoses, both in otherwise unperturbed intestines and in damaged ones (Fig. 5). Consistently, hypoxia also inhibited tumor growth, tracheal coverage and mitoses (Fig. 8). Since oxygen is necessary for metabolism and mitosis, and hypoxia limits the amount of oxygen available to be transported to the ISCs, this could explain the inability of the extra branches to support regenerative and tumor growth in the midgut in hypoxic conditions (Fig. 8). In summary, we propose that intestinal tissue homeostasis is maintained by oxygen availability. Both intestinal regenerative responses and growth of ISC-derived tumors depend on the levels of oxygen reaching the midgut epithelium through the tracheal branches. In addition, tracheal branching is dynamically regulated by ROS- and oxygen-dependent signaling emanating from the epithelium itself. Thus, a feed-forward loop of ROS inducing tracheal branching increases oxygen supply to the epithelium. We suggest that increased oxygen availability potentially produces more ROS that drive ISC mitosis via activation of key signaling pathways, such as Jak/Stat and EGFR (Fig. 8l).

## Materials and Methods

### Fly stocks and maintenance

*Drosophila* stocks were maintained at 18^°^C on a 12:12 hour light:dark cycle on standard fly food containing propionic acid (Sigma) and Tegosept (Genesee).

The following Gal4 lines were used for tissue and cell-type specific expression: *yw*; *btl-Gal4 UAS-srcGFP* ^8^, *w; btl-Gal4 UAS-srcGFP tub-Gal80^ts^/CyO* (this work), *w; esg-Gal4^NP5130^ UAS-src*GFP ^8^, *w; esg-Gal4 UAS-GFP tub-Gal80^ts^* ^21^, *w; Su(H)-Gal4 UAS-CD8-GFP tub-Gal80^ts20^/CyO* (this work; *Su(H)-Gal4* was a gift from S. Hou ^23^), *Myo1A-Gal4 UAS-GFP/CyO Su(H)-Gal80 tub-Gal80^ts^/TM6C* (this work), *w; Myo1A-Gal4 UAS-mcherry tub-Gal80^ts^/CyO* (this work), *w; trh-Gal4^14D03^* (BDSC# 47463), *dSRF-Gal4* (BDSC# 25753); *w; dSRF-Gal4 UAS-GFP* ^96^, *w; esg-Gal4 UAS-GFP tub-Gal80^ts^/CyO; QF6 QUAS-mtdTomato/TM6B* (this work; QF6 was a gift from C. Potter ^52^), *w; Myo1A-Gal4 UAS-nls-mCherry tub-Gal80^ts^; QF6 QUAS-mCD8GFP* (this work; ^52^), *w; btl-Gal4 UAS-src-GFP tubGal80^ts^; sima^KG^/T(2;3)* (this work), *w; Myo1A-Gal4 UAS-nls-mCherry tub-Gal80^ts^; sima^KG^/T(2;3)* (this work), *w; esg-Gal4 UAS-eGFP tub-Gal80^ts^/CyO; UAS-Ras1A^Q13^/TM6C* (this work); w; *ppk-Gal4/ CyO* (BDSC# 32078), w; *UAS-CD8::GFP/II* (BDSC# 5137), *w; mira-GFP/CyO; trh-Gal4^14D03^-Gal4/TM6B* (this work, *mira-GFP* was a gift from A. Bardin ^97^).

The following UAS strains were used: *UAS-btl^DN^* ^3^, *UAS-λbtl/TM3,Sb* ^3^, *UAS-λbtl* (BDSC# 29046), *UAS-bnl* (BDSC# 64231), *UAS-sima^V10,RNAi^(II)* (BDSC# 26207), *UAS-simaB* (BDSC# 9582) ^63^, *UAS-Cat^RNAi^* (VDRC# 6283), *UAS-Sod1A* (BDSC# 24750), *UAS-Cat* (BDSC# 24621), *UAS-bnl^RNAi1^* (VDRC# 101377), *UAS-bnl^RNAi2^* (VDRC# 5730), *UAS-bnl^RNAi^ (BDSC# 34572), UAS-Ras^V12^* (BDSC# 4847) and *UAS-Notch^DN^* ^98^, *UAS-Notch^RNAi^* (BDSC# 7077), *UAS-Duox^RNAi^* ^99^.

The following mutants and reporter lines were used: *w*^1118^ (BDSC# 3605), *sima^KG07607^* (BDSC #14640, isogenized in CP lab), *btl*^MI03286^ (BDSC# 36229), *btl^f02864 100^*, *bnl^MI01635^* (BDSC# 34228), *bnl^e03115 100^*, *w; UAS-src-GFP; bnl-Gal4^NP2211^* (this work, originating from BDSC# 112825), *w; ldh-Gal4 UAS-nls-GFP* and *w; ldh-Gal4 UAS-src-GFP* (this work, *ldh-Gal4* was a gift from P. Wappner ^63^), *esg-lacZ*^606^ (BDSC# 10359), *Dl-lacZ*^05151^ (BDSC# 11651) and *Su(H)-lacZ* (BDSC# 83352).

For crosses involving driver lines with “*tubulin-Gal4, Gal80^ts^*” were set up at 18^°^C, F1 adult progeny were collected within two days of emerging from the pupae. These flies were transferred to 29^°^C for the expression of the transgene including *RNAi* construct of interest. Flies were used within 10 days of emerging for all the stress experiments. Flies were transferred to new food every 2 days except for weekends. Fly stocks used for the experiments are listed in the reagent table as a supplementary information.

Control and Ras^V12^ MARCM clones ^101^ were generated by crossing *yw hs-FLP tub-Gal4 UAS-nls-EGFP; FRT82B tubGal80* ^102^ to *w; FRT82B arm-lacZ* (BDSC# 7369) and *w; UAS-Ras^V12^; FRT82B arm-lacZ* (this work), respectively. In short, 3-day-old flies reared at 25°C were heat shocked at 37°C in a circulating water bath for 60 min, once a day for 2 sequential days. Then, they were maintained at 25°C for one week until dissection.

### Oral administration of bacteria and chemicals

Female mated mature flies were used for all feeding assays.

#### P. aeruginosa

*P.a.* feeding was performed as previously described ^33^. Briefly, a single colony from the *P. aeruginosa* strain PA14 was grown at 37°C to OD_600nm_=3, corresponding to 5×10^9^ bacteria/ml. Female mature flies of the desired genotype were starved for 5 hours and added in groups of 10 per fly vial containing a cotton ball at the bottom impregnated with 5 ml of 0.5ml PA14 OD_600nm_=3, 1ml 20% sucrose and 3.5ml dH_2_O. For uninfected control, 5ml of 1ml sucrose 20% and 4ml dH_2_O was used. Flies were incubated for 48 hours at 25°C or 29°C (for all experiments utilizing the Gal4-UAS system, unless otherwise noted).

#### P. entomophila

*P.e*. was cultured in LB with rifampicin (100µg/ ml, Sigma-Aldrich, USA) at 29^°^C. Initial 50 ml overnight culture was transferred to 150 ml media for further 24 hours at the same temperature. *P.e.* was spun-down, mixed with 5% sucrose, yeast-extract and bacterial supernatant. Flies starved for 4 hours with water were fed with this *P.e.* mix. 5% sucrose was fed to the control groups at the same time.

#### H_2_O_2_

H_2_O_2_ feeding was performed as previously described^103^. 30 % H_2_O_2_ solution (EMSURE) was stored at 4°C. 500 μl of 1% H_2_O_2_ with 4% sucrose solution was pipetted onto a sterile cotton ball at the bottom of an empty vial, and 4% sucrose solution without H_2_O_2_ was used as control. Flies were transferred to treatment vials and incubated for 48 hours at 25°C before dissection, unless otherwise noted.

#### Paraquat

This treatment was performed as described previously^104, 105^. PQ powder (Sigma) was diluted in ddH_2_O and stored as a stock of 100mM at 4°C. A 10mM PQ solution prepared fresh in 4% sucrose was used as test food and 4% sucrose solution without PQ as a control. Flies were transferred to the empty fly food vial containing only one 23mm Whatman paper disc soaked with 250μl of test or control food and incubated for 24 hours at 25°C, unless otherwise noted.

#### *P. aeruginosa* mutants supernatant administration (MvfR, PhzS, PhzM)

The mutant strains MvfR, PhzS and PhzM were taken from a *P.a.* library designed by F. Ausubel^69^. Cell-free supernatants of PA14, MvfR, PhzS and PhzM were prepared by centrifugation and filtering of the supernatant with 0.2 μm filters. Fresh cell-free supernatant solutions were prepared daily for 2 days. Flies were transferred to the vials with the wet cotton balls impregnated with 5 ml of a solution composed of 80% LB medium or supernatant and 20% sucrose.

### Dissections and Immunohistochemistry

Adult midgut dissections and immunohistochemistry were performed as previously described ^33^. Following dissections on Sylgard (VWR) plates in 1x PBS (130mM NaCl, 70mM Na_2_HPO_4_, 30mM NaH_2_PO_4_). Dissected midguts were fixed in 4% Formaldehyde (Polysciences) in 1x PBS for 30 minutes at room temperature. After 3 rinses with PBS, the tissues were incubated in blocking PBT solution (1x PBS, 0.5% BSA, 0.2% TritonX-100) at room temperature for at least 20 minutes. Addition of the primary antibody in PBT was followed by incubation for 16 hours at 4°C. Primary antibody was washed 3 times for 10 min at room temperature with PT (1x PBS, 0.2% TritonX-100). Secondary antibodies and DAPI (Sigma) are added for 1-2 hours at room temperature. Excess antibodies are washed in PT (as before) and the midguts are mounted in Vectashield (Vector). Primary antibodies: rabbit-anti-pH3 (Millipore 1:4000), rabbit anti-cleaved-Caspase-3 (#9661 Cell Signaling Technology 1:400), mouse-anti-Prospero (DSHB 1:100), mouse anti-β-gal (Promega 1:500), chicken-anti-GFP (Invitrogen 1:1000), rabbit anti-GFP (1:3000; Invitrogen) DAPI (Sigma, 1:3000 of 10mg/ml stock) (staining of the nuclei). Secondary antibodies against mouse, rabbit or chicken conjugated to Alexa Fluor 488 and 555 (Invitrogen) were used at 1:1000.

### ROS detection

For ROS detection, the Dihydroethidium (DHE) staining method was used. Midguts were dissected in 1x PBS and incubated in 30μM DHE (Invitrogen), for 5 minutes in the dark, at 37°C. DHE was washed 3 times with 1x PBS. The guts were mounted and immediately imaged by confocal microscopy.

### RT-qPCR

Dissected midguts were collected in Qiazol (Qiagen). Total RNA was extracted from 20 midguts per strain per condition per biological replicate using the Qiazol protocol. 800 ng of total RNA were freed from genomic DNA using the RQ1 RNase-Free DNase Kit (Promega). Reverse transcription was performed using 145,4ng of the total DNase-treated RNA using the TaKaRa PrimeScript RT Master Mix Kit. qPCR amplification was performed using gene specific primers with the following amplification program: 95°C for 30 seconds (initial denaturation), 40 cycles of 95°C for 10 seconds (denaturation), 60°C for 30 seconds (annealing), 65°C for 30 seconds (extension) and 65°C for 1 minute (final extension). Primer sequences for each gene are shown in Table 1. The expression of the genes of interest was normalized to the expression levels of two reference genes, *rpl32* and *gapdh1* using the 2-ΔΔCt method. Data were analyzed using the Bio-Rad CFX Manager 3.1 program and graphs were prepared in Excel. For all experiments at least 3 biological replicates were used to calculate the average and the standard deviation.

The primer pairs used for qPCR analyses are the following:

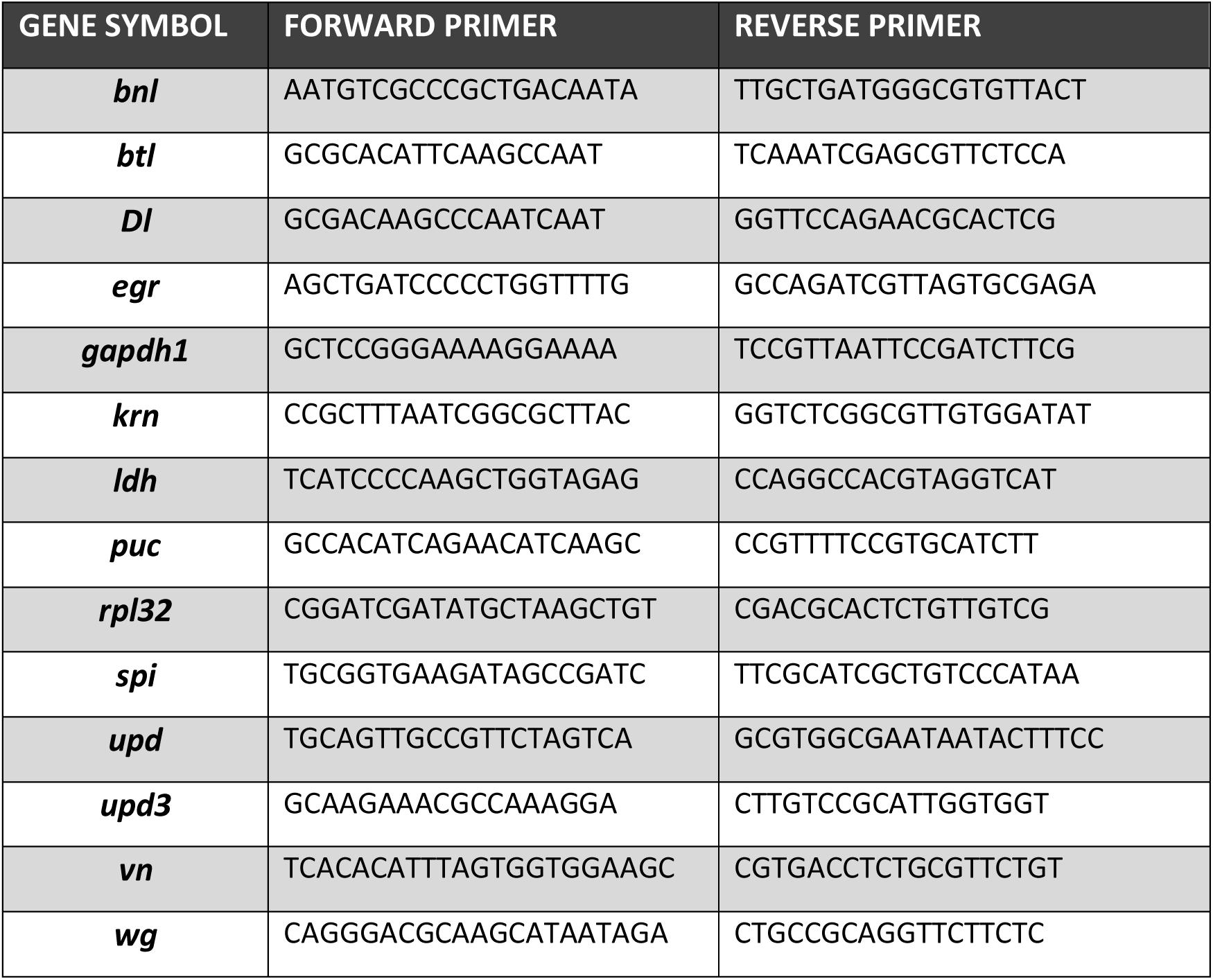

### Midgut Tumorigenesis

Tumors (*Ras^V12^*, *Ras^Q13^*, *Notch^DN^*, *Ras^V12^ Notch^RNAi^*) were induced in the midgut intestinal progenitor population via the Gal4-UAS-Gal80^ts^ system^50, 74^. Tumor-generating crosses were reared at 18°C, and emerging adult progeny were maintained at 18°C to mature for 5-7 days. After maturation, mated adult females were transferred to 29°C to induce transgenes. For *Ras^V12^* tumors, flies were induced for 1, 3 and 5 days, for *Notch^DN^* tumors flies were induced for 4, 7 and 10 days. For *Ras^Q13^* tumors flies were induced for 1 and 2 days. For *Ras^V12^ Notch^RNAi^* tumors, flies were induced for 7 days.

For MARCM tumors, the crosses were reared at 25°C and mature 3-5 days old adult female progeny with the correct genotype were heat-shocked for 1 hour at 37°C in a water bath. A time-course analysis (3, 7, 10, 14 days) was performed to identify the time point when large clones were readily visible. Large close were obtained 14 days after heat shock. Images were acquired in a Zeiss Axioscope A1 to show GFP fluorescence of the clone and bright-field image of the trachea.

### Survival assay

Flies of the appropriate genotypes were collected within 1-2 days after hatching at 18^°^C. The flies were transferred to 29^°^C and the number of dead flies were counted every 2 days while changing the food. The Kaplan-Meier log-rank test was used to assess statistical significance and the data were plotted using Graphpad Prism 9.

### Tumor transplants

Midguts bearing control progenitor cells and tumors were dissected out and cut into pieces. Cut pieces were aspirated into a hypodermic syringe and carefully injected into the abdomen of the anesthetized recipient flies. Recipient flies with transplanted midgut pieces were incubated at 29^°^C until observation was complete. The percentage of tumor growth was calculated by counting the number of flies bearing tumors compared to the number of flies injected with the tumor pieces. The percentage of distal colonization was calculated by counting the number of tumor-bearing flies showing tumors above the thorax region (including the head) away for the site of injection in the abdomen to the number of flies injected with the tumor pieces.

### Hypoxia

*Drosophila* strains were reared in normoxic (21% O_2_) and hypoxic conditions (5% O_2_) for 12 hours. For the hypoxic conditions the Modular Incubator Chamber (MIC-101, Billups – Rothenberg Inc.), was used connected to a cylinder that contained a mixture of gasses with 4% O_2_. The flies for both normoxic and hypoxic conditions were transferred into new vials containing the oral experimental condition and were covered with net (instead of cotton). The chamber was moved into a 29°C incubator overnight (12 hrs). For all the combined hypoxia-infected and hypoxia-tumor conditions, hypoxia was the last step, meaning that the flies before dissection were reared in low oxygen tension. The flies that were reared in hypoxia were dissected first to reduce contact with the atmospheric oxygen.

### Image acquisition and analysis

Full midgut images (as those used in Extended Data Fig. 1 and 3) were acquired with a Leica MZ16F Fluorescent Stereomicroscope equipped with GFP and DAPI filters.

Stacks of optical sections were acquired using the Leica TCS SP2 DMIRE2 Scanning Confocal Microscope using a 40x oil objective (zoom 1x, average 4). Images to be compared were acquired using the exact same settings. Bright-field images were acquired using the fluorescent microscope Zeis Axioscope A.1 at 20x magnification.

Measurements of tracheal length were performed using the NeuronJ Plugin of ImageJ 1.34s software (*ImageJ – Plugins – NeuronJ. File – Open image file*). The monitoring of the tracheal system can take place in two independent ways. The first relies on the quantification of the fluorescence intensity of the trachea and the other on the quantification of the tracheal branches. For trachea quantification, images were taken using a Leica TCS SP2 DMIRE2 confocal microscope, at 40Χ magnification, zoom1, image size: 1024×1024, 0.5μm step size, sequential scan mode DAPI/RFP, ∼18-20 serial sections for both the anterior and the posterior areas of the midgut. Importantly, to achieve accurate comparisons, all the images were acquired under the same GFP laser settings. Numerous intestines were photographed from the anterior (R2 region), the posterior (R5) region and occasionally from the middle (R3, R4a) midgut (specified throughout the text). Using the NeuronJ tool (*Add tracings*), an ImageJ Plugin for Neurite Tracing and Analysis found in the ICY software (http://icy.bioimageanalysis.org/), we highlighted each tracheal branch along on the surface of the midgut. We avoided highlighting the large, thick external tracheal vessels that do not adhere to the gut surface. Then, we summed the full length of the branches (*Measure tracings*). Normalization to the surface of the midgut was performed for each image before plotting. We tested this image quantification method and compared it to other methods. We found that this was the most accurate method for intestinal tracheal measurements, and thus, we used it for all subsequent experiments.

Graphs were generated using Graphpad Prism 8 by calculating the average and standard deviation values. Bright-field images from the Zeiss Axioscope A.1 were also used to measure the tracheal length using the NeuronJ Plugin. In the Edgar lab, the images of the midguts with trachea (Fig. 1e, f, Fig. 7b) were taken at the same magnification with a 40x water objective using a Leica SP8 confocal microscope. Bright-field DIC images (Fig. 1k, Fig. 7a,c) were taken with a Leica DM microscope and grouped anonymously for quantifying the phenotypes. The tracheal branches were categorized into primary, secondary, and tertiary branches according to their thickness with thickest being the primary trachea. No significant differences in the number of primary or secondary trachea were observed across the groups. The number of tertiary tracheal branches per field was counted manually from the images.

The numbers of pH3-positive cells were counted under the fluorescent microscope (Zeiss Axioscope A.1 and Nikon Eclipse Ti) at 20x magnification along the whole midgut. Graphs were generated using Graphpad Prism 9 by calculating the average and standard deviation values.

## Data availability

The authors declare that the data supporting the findings of this study are available within the paper and its supplementary information files.

## Acknowledgements

The authors would like to thank the BDSC, VDRC and the TRiP for fly stocks; Allison Bardin, Steven Hou, Jordi Casanova, Thomas Kornberg, Chris Potter, Pablo Wappner and Anastasia Ignatiou for fly stocks; Eric Snyder for the Nikon Ti microscope usage; the DSHB for antibodies. This project was supported by FP7-PEOPLE-2011-CIG-303727, the Fondation Santé and the Cyprus RPF-EXCELLENCE/0918/0082 to C.P., and by ERC AdG 268515, DFG SFB873, the Huntsman Cancer Foundation and NIH GM124434 to B.A.E.

## Author contributions

V.T., M.M.R. and M.P. designed and performed experiments, and analyzed data. I.C. and K.K. performed experiments and analyzed data. H.M., V.B. and B.I. assisted with experiment execution. B.A.E. and C.P. conceived and supervised the project. C.P. compiled the data and wrote the manuscript with the help of M.M.R., V.T. and B.A.E.

## Figure legends

**Extended Data Figure 1.**
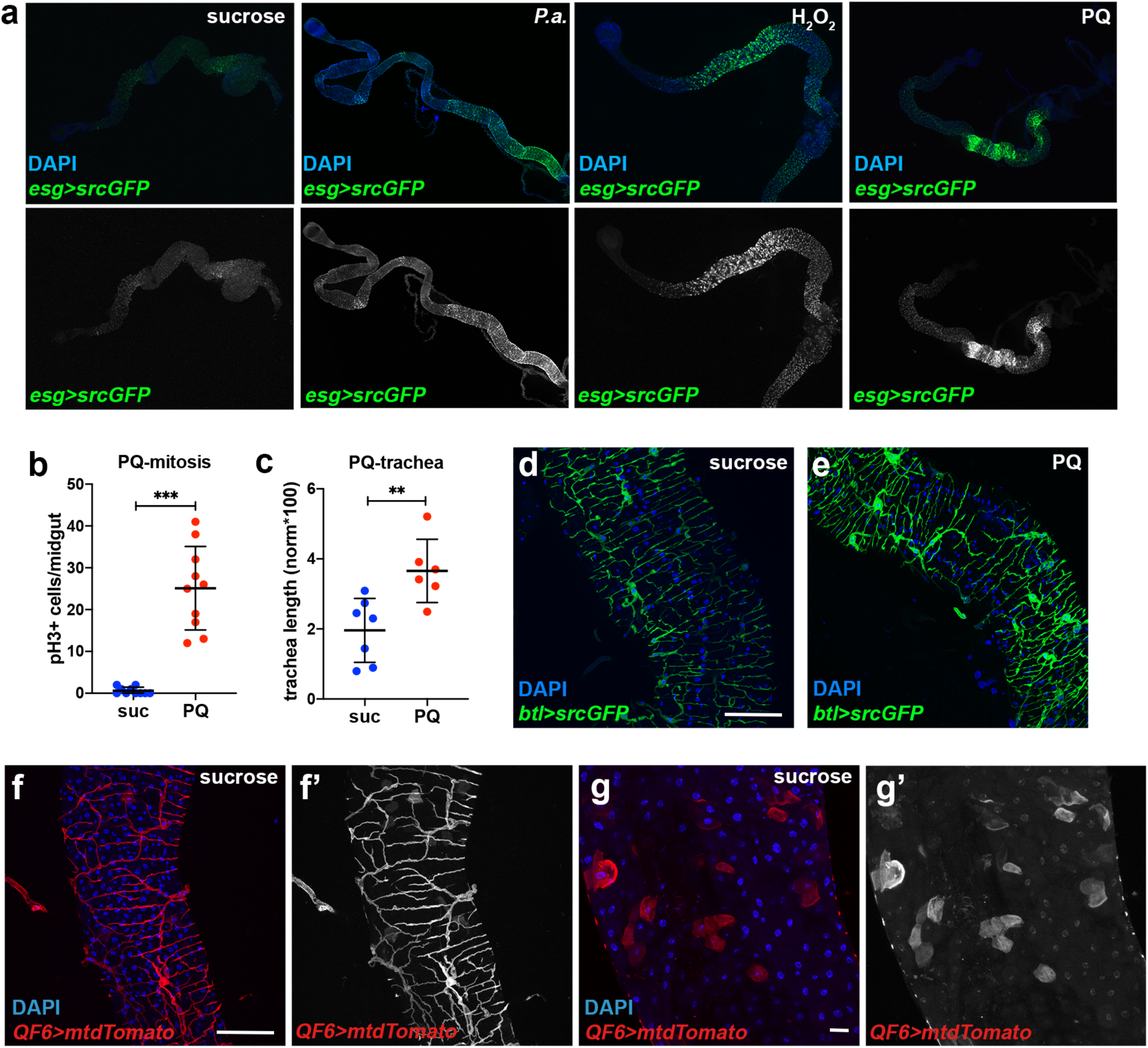
Infection and oxidative damage increase *esg>GFP+* cells in the midgut and associate with increased TTC branching. **a.** Adult midgut intestinal progenitors labeled with *esg^NP5130^*–*Gal4>UAS-srcGFP* were imaged in unchallenged conditions (4% sucrose) and upon oral *P.a.* infection (48hrs), and feeding with H_2_O_2_ (48hrs) and PQ (24hrs). DAPI (blue) in the upper panels stains all midgut nuclei. The bottom panels show the GFP-labeled progenitors separately. *P.a.* and PQ expanded the intestinal progenitors with a posterior midgut bias, whereas H_2_O_2_ exhibited an anterior midgut bias. **b-c.** Quantification of midgut mitosis (b) and TTC branching (c) in PQ-treated flies. **d-e.** Posterior midgut (R4) images of *btl-Gal4>UAS-srcGFP* flies in baseline conditions (sucrose) and upon PQ feeding. DAPI (blue) staining all the nuclei. Single channel images of the GFP are shown in d’-e’. **f-g.** Posterior midgut images of *QF6>QUAS-mtdTomato* flies in baseline conditions exhibit tracheal expression of the reporter. Midgut epithelial ECs with low expression of the reporter are visible is zoomed image (g). Single channel images of the Tomato are shown in f’-g’. Scale bars: 75 μm in d-g. Error bars correspond to standard deviation. Statistical significance (t-tested for b, and U-tested for c): ns, not significant, * 0.01<*p*≤0.05, ** 0.001<*p*≤0.01 and *** *p*≤0.001.

**Extended Data Figure 2.**
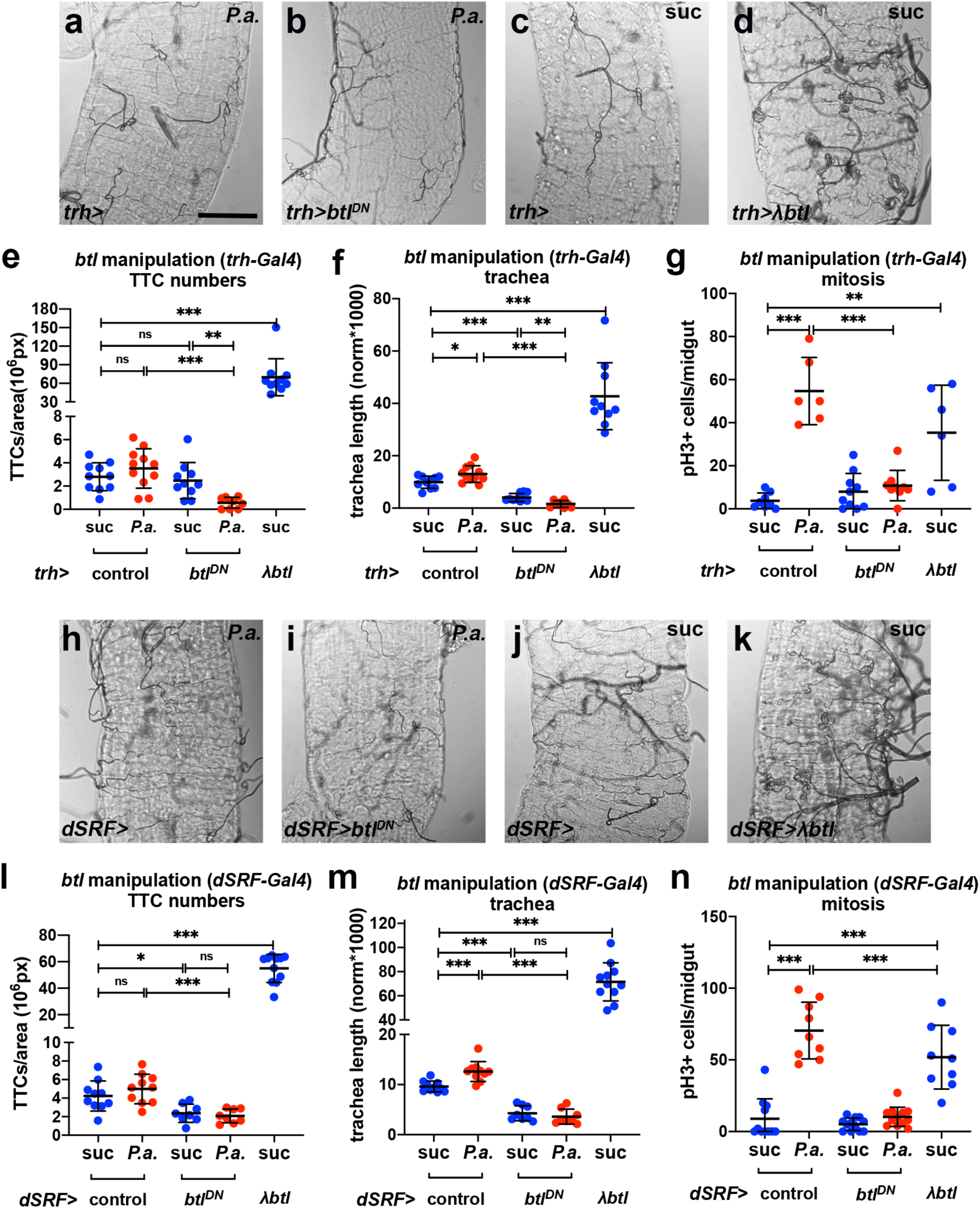
The FGFR/Btl is necessary and sufficient for midgut TTC branching and ISC mitosis. **a-b.** Brightfield images of the tracheae of *P.a.* infected R5 regions of the midgut in *trh-Gal4* control and *trh-Gal4*>*UAS-btl^DN^*-expressing flies. **c-d.** Brightfield images of the tracheae of uninfected R5 regions of the midgut in *trh-Gal4* control and *trh-Gal4*>*UAS-λbtl*-expressing flies. **e-g.** Quantification of TTCs (e), TTC branching (f), and midgut mitosis (g) upon *trh-Gal4*-driven *btl* manipulation with or without *P.a*. infection. **h-k.** Brightfield images of the tracheae of *P.a.*-infected R5 regions of the midgut in *dSRF-Gal4* (h) control and *dSRF-Gal4*>*UAS-btl^DN^* (i) as well as control (j) and *dSRF-Gal4*>*UAS-λbtl* (k)-expressing flies. **l-n.** Quantification of TTCs (l), TTC branching (m), and midgut mitosis (n) upon *dSRF-Gal4*-driven *btl* manipulation with or without *P.a*. infection. Scale bars: 75 μm. Error bars correspond to standard deviation. Statistical significance (t-tested): ns, not significant, * 0.01<*p*≤0.05, ** 0.001<*p*≤0.01 and \*\*\**p*≤0.001.

**Extended Data Figure 3.**
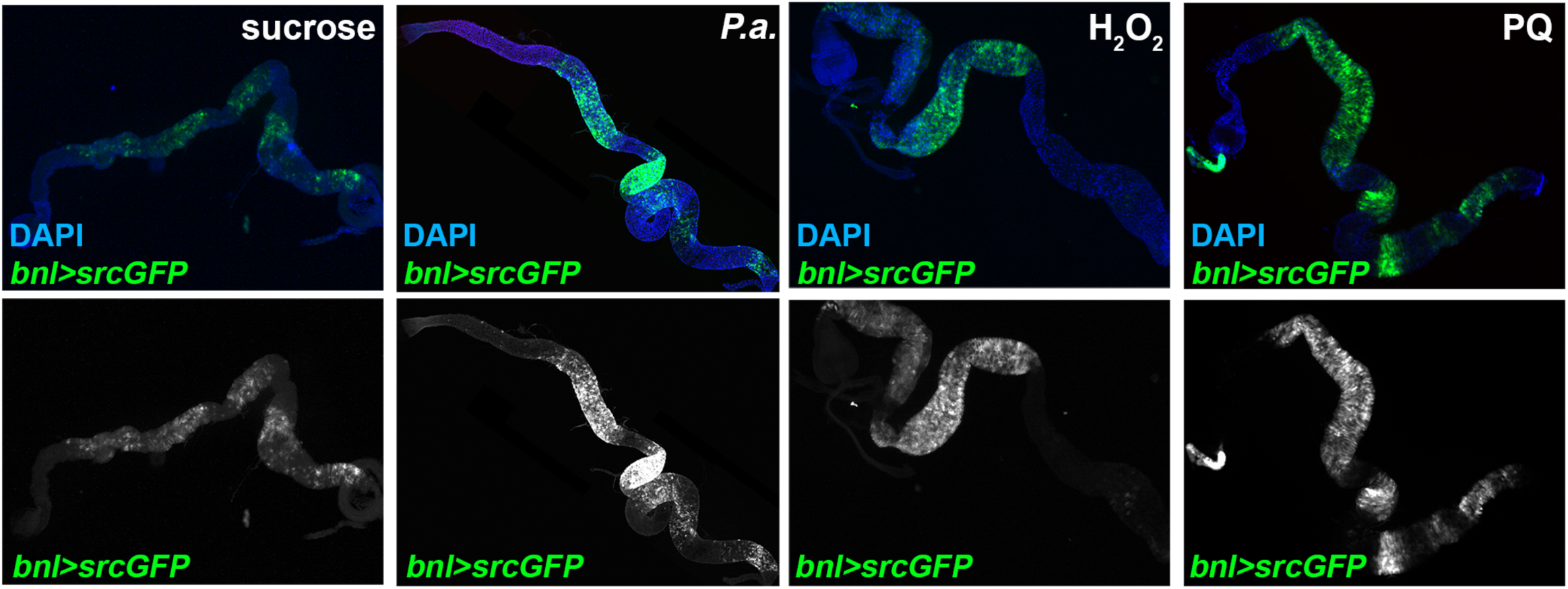
Infection and oxidative damage induce *FGF/bnl* in the midgut epithelium. Adult midgut *bnl*-expressing cells labeled with the reporter *bnl-Gal4>UAS-srcGFP* were imaged in unchallenged conditions (4% sucrose) and upon oral *P.a.* infection (48hrs), feeding with H_2_O_2_ (48hrs) and PQ (24hrs). DAPI (blue) in the upper panels stained all midgut nuclei. The bottom panels show the GFP-labeled *bnl* expressing cells separately. *P.a.* and PQ induced the reporter throughout the midgut, whereas H_2_O_2_ exhibited an anterior midgut bias.

**Extended Data Figure 4.**
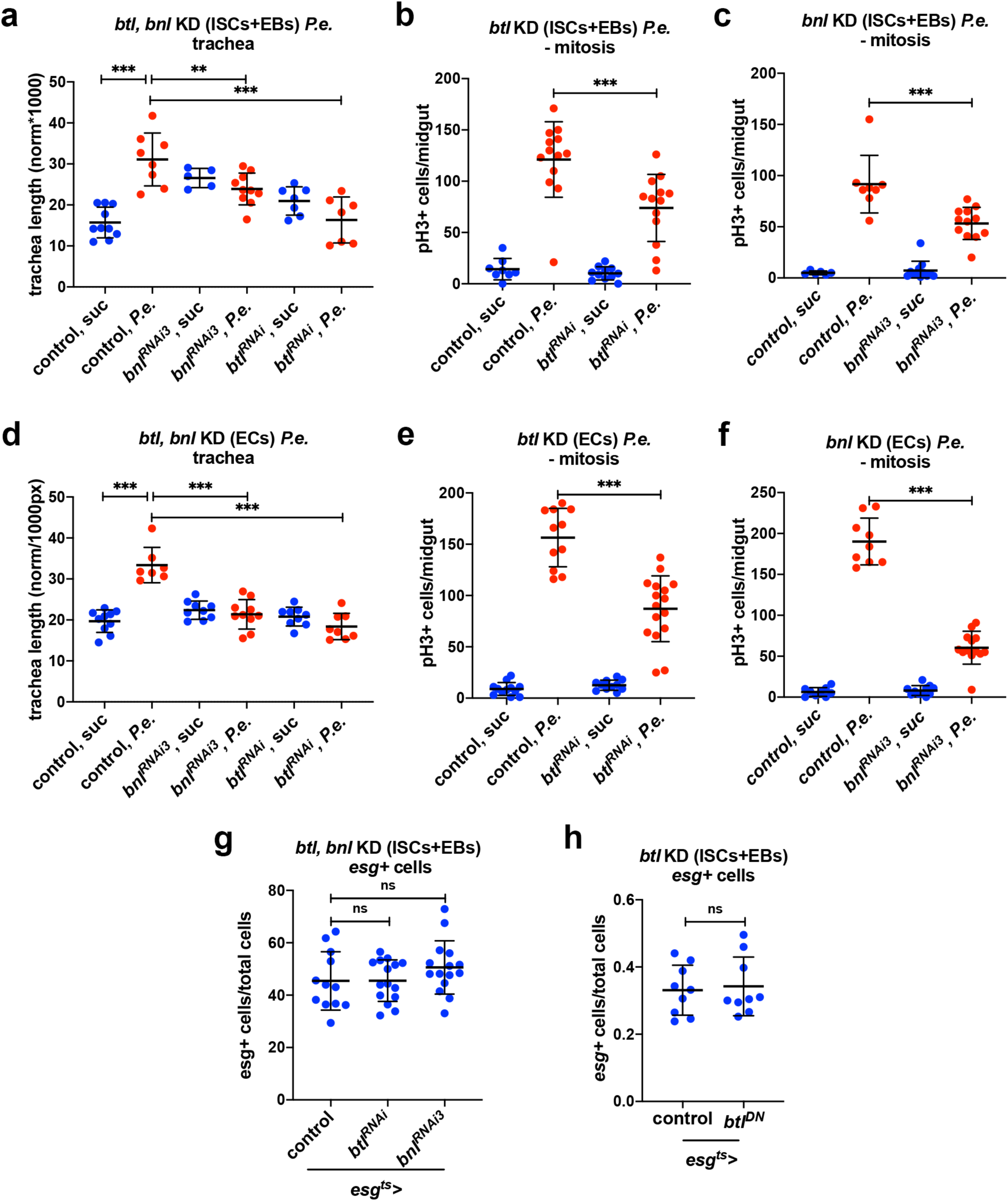
Btl/Bnl signaling in the epithelial cells is necessary for efficient tracheal remodeling and mitosis in response to infection. **a-c.** Quantification of TTC branching (a) and midgut mitosis (b-c) upon progenitor-specific knockdown of *btl* (*btl^RNAi^*) and *bnl* (*bnl^RNAi3^*) with or without *P.e.* infection **d-f**. Quantification of TTC branching (d) and midgut mitosis (e-f) upon EC-specific knockdown of *btl* (*btl^RNAi^*) and *bnl* (*bnl^RNAi3^*) with or without *P.e.* infection. **g.** Quantification of *esg*+ progenitors as a percent of total number of cells in the posterior regions of the midgut upon progenitor-specific knockdown of *btl* (*btl^RNAi^*) and *bnl* (*bnl^RNAi3^*). **h.** Quantification of *esg*+ progenitor cells/total number of cells in the posterior midgut upon progenitor-specific knockdown of *btl* (*btl^DN^*). Error bars correspond to standard deviation. Statistical significance (t-tested): ns, not significant, * 0.01<*p*≤0.05, ** 0.001<*p*≤0.01 and *** *p*≤0.001.

**Extended Data Figure 5.**
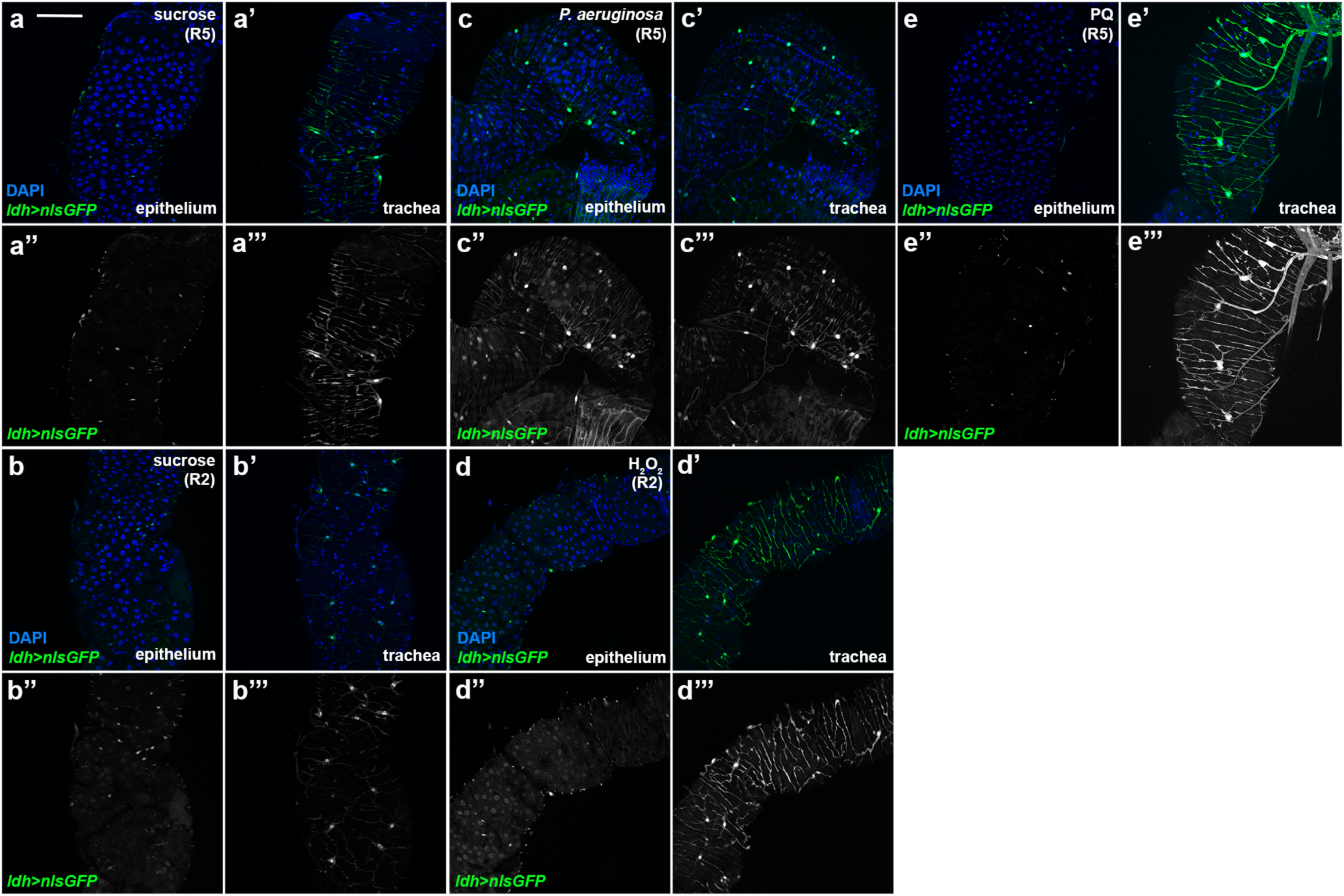
Infection and oxidative damage activate Hif-1α/Sima in the midgut epithelium and the visceral TTCs. Hif-1α/Sima activation was monitored via the *ldh-Gal4>UAS-nlsGFP* reporter expression in the adult midgut epithelium and the intestinal trachea of the R5 region in unchallenged flies (sucrose) and upon *P.a.* and PQ treatment (a, c, e), and of the R2 region in unchallenged flies (sucrose) and upon H_2_O_2_ feeding (b, d). Epithelial sections (a-d) and trachea surface sections (a’-d’) of the same midguts were imaged. DAPI (blue) in a-d and a’-d’ stains all the nuclei. a’’-d’’ and a’’’-d’’’ correspond to separated channels for reporter expression in the epithelium and the intestinal trachea, respectively. The *ldh-Gal4>UAS-nlsGFP* reporter was expressed in cells of the midgut epithelium and in the midgut TTCs in baseline conditions in the anterior (R2 in b, b’) and posterior (R5 in a, a’) midgut. *P.a.* (c, c’), H_2_O_2_ (d, d’) and PQ (e, e’) induced the reporter in the epithelium and the trachea at varying degrees. All images were acquired at the same confocal settings as their respective controls. Scale bars: 75 μm.

**Extended Data Figure 6.**
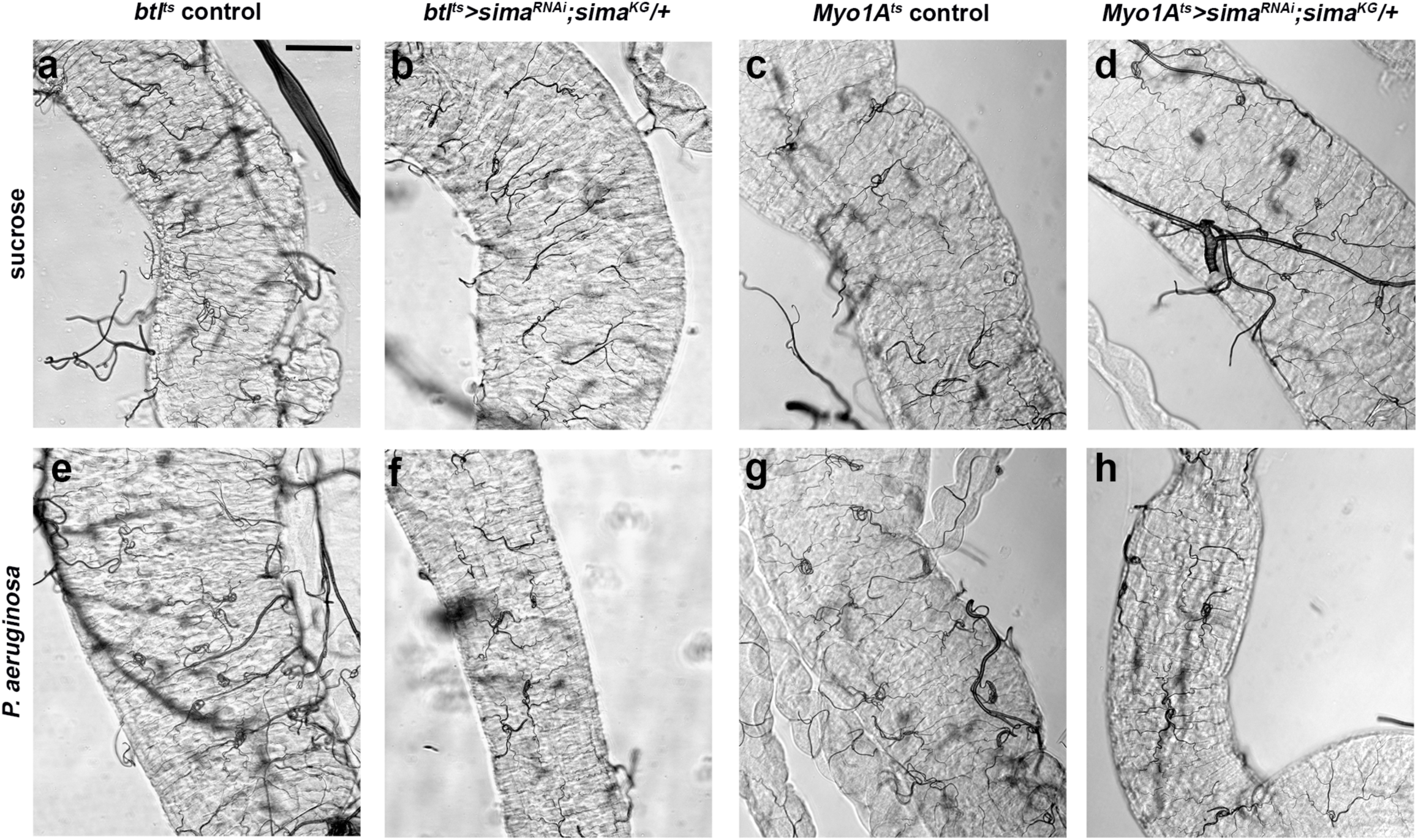
Hif-1a/Sima is necessary in the intestinal epithelium and the trachea for TTC branching. **a-b.** Brightfield images of the midgut TTCs (R5 region) upon trachea-specific (via *btl-Gal4*) *sima* knockdown in the background of heterozygous *simaKG* in baseline conditions. **c-d.** Bright-field images of the midgut TTCs (R5 region) upon EC-specific (via *Myo1A-Gal4*) *sima* knockdown in the background of heterozygous *simaKG* in baseline conditions. **e-f.** Bright-field images of the midgut TTCs (R5 region) upon trachea-specific (via *btl-Gal4*) *sima* knockdown in the background of heterozygous *sima^KG^* in *P.a.-*infected conditions. g-h. Bright-field images of the midgut TTCs (R5 region) upon EC-specific (via *Myo1A-Gal4*) *sima* knockdown in the background of heterozygous *sima^KG^* in *P.a.-*infected conditions. The images correspond to examples of those quantified for Fig. 4e,i. Scale bars: 75 μm.

**Extended Data Figure 7.**
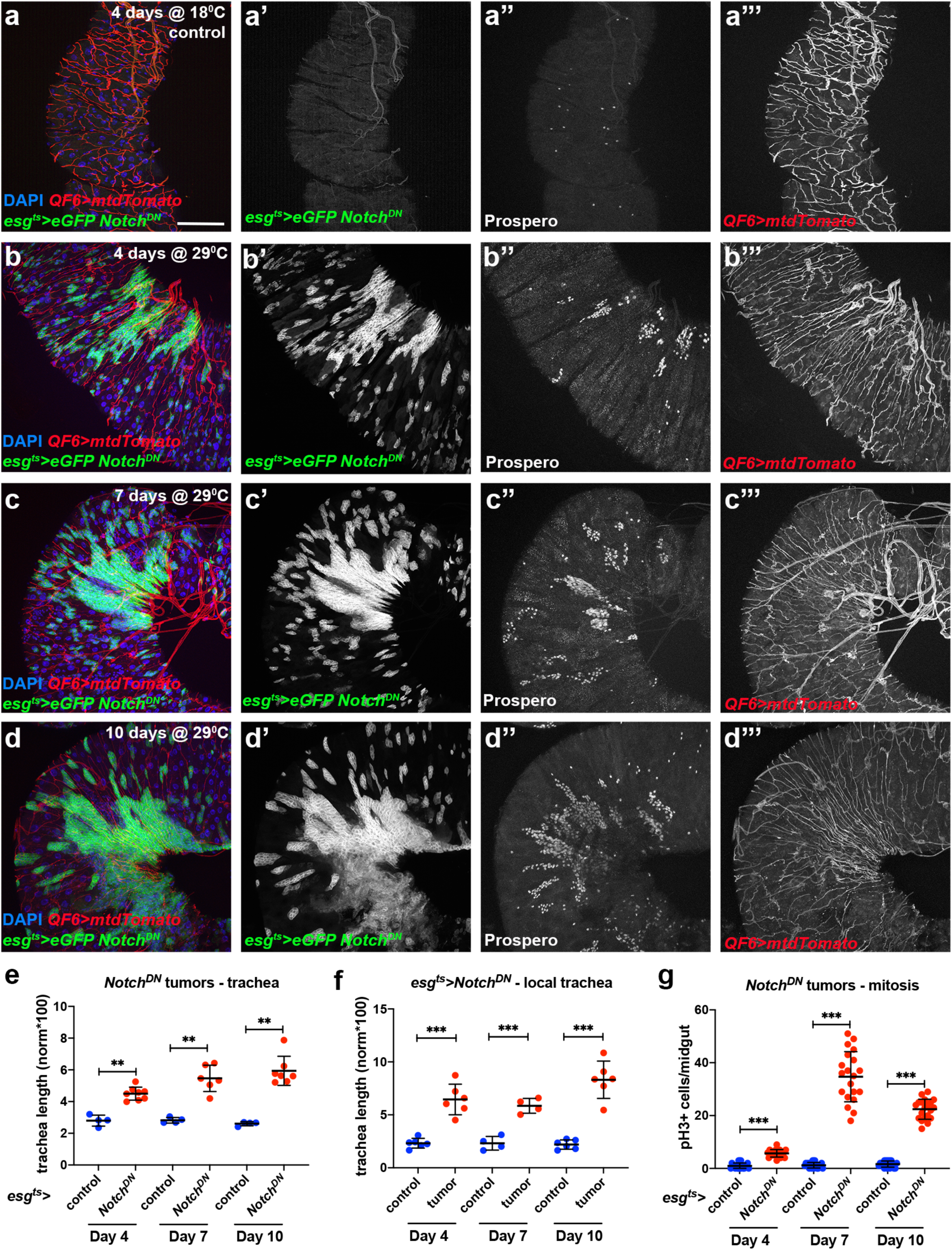
Time-course analysis of *Notch^DN^* progenitor-derived midgut tumors. **a-d.** The R4a region of control (reared for 4 days at 18^°^C) (a) and tumorous midguts (reared for 4, 7 and 10 days at 29^°^C) (b-d) of the *esg-Gal4 UAS-eGFP tub-Gal80^ts^>UAS-Notch^DN^* genotype with concomitant expression of *QF6>QUAS-mtdTomato* (red) to label the trachea. DAPI (blue in a-d) is used to label all midgut nuclei and Prospero (a’’-d’’) labels the EEs. a’-d’, a’’-d’’ and a’’’-d’’’ correspond to the individual channels for eGFP, Prospero and Tomato-labeled trachea, respectively. Scale bars: 75 μm. **e-g**. Quantification of TTC branching in the R4a of control (*Notch^DN^* uninduced) and *Notch^DN^*-expressing midguts (e), in the *Notch^DN^* tumor-region vs. neighboring non-tumor area on the same image (f), and midgut mitosis of control (*Notch^DN^* uninduced) and *Notch^DN^*-expressing midguts (g) during a time-course analysis at 4, 7, and 10 days post-tumor induction. Scale bars: 75 μm. Error bars correspond to standard deviation. Statistical significance (t-tested): ** 0.001<*p*≤0.01 and *** *p*≤0.001.

**Extended Data Figure 8.**
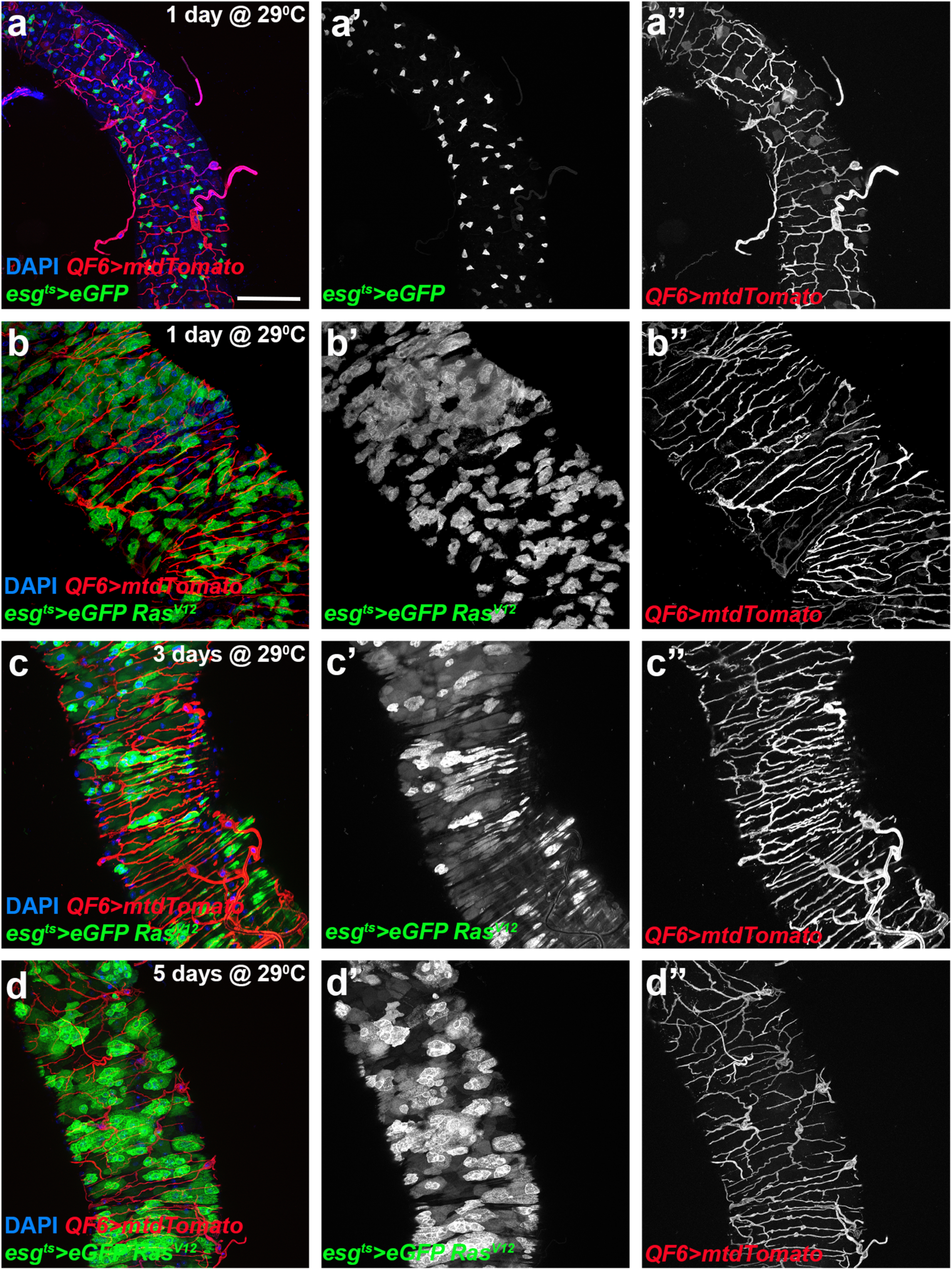
Time-course analysis of *Ras^V12^* progenitor-derived midgut tumors. **a-d.** The R5 region of control *esg-Gal4 UAS-eGFP tub-Gal80^ts^* (reared for 1 day at 29^0C^) and *esg-Gal4 UAS-eGFP tub-Gal80^ts^*>*UAS-Ras^V12^*-tumor bearing midguts (reared for 1, 3 and 5 days at 29^°^C) with concomitant expression of *QF6>QUAS-mtdTomato* (red) to label the trachea. DAPI (blue in a-d) was used to label all midgut nuclei. a’-d’ and a’’-d’’ correspond to the individual channels for the eGFP and the Tomato-labeled trachea, respectively. Scale bars: 75 μm.

**Extended Data Figure 9.**
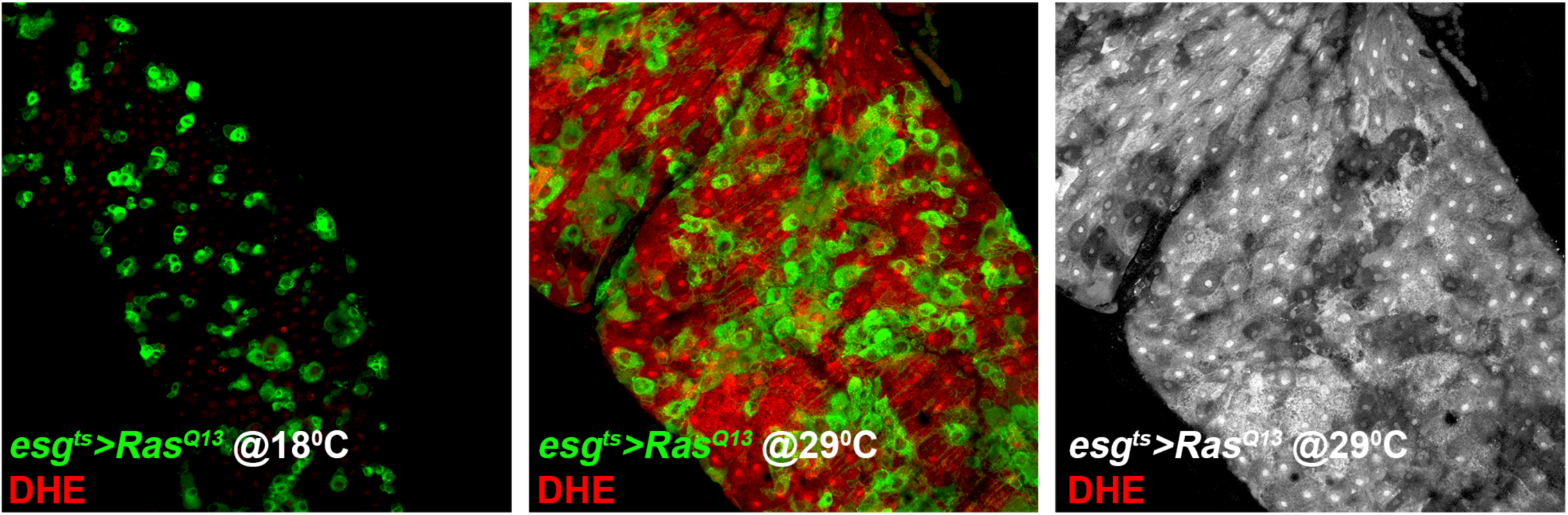
Increased ROS levels in the midgut with *Ras^V12^* progenitor-derived tumors. The R5 region of *esg-Gal4 UAS-eGFP tub-Gal80^ts^*>*UAS-Ras^v12^*-tumor bearing midguts from flies reared at 18^°^C and 29^°^C with DHE (red) co-staining for ROS levels. The flies were kept @18^°^C and 29^°^C for 3 days. The right-most panel shows the channel for DHE staining only.

## References

1. Ghabrial, A., Luschnig, S., Metzstein, M.M. & Krasnow, M.A. Branching morphogenesis of the Drosophila tracheal system. Annu Rev Cell Dev Biol 19, 623–647 (2003).

2. Hayashi, S. & Kondo, T. Development and Function of the Drosophila Tracheal System. Genetics 209, 367–380 (2018).

3. Sato, M. & Kornberg, T.B. FGF is an essential mitogen and chemoattractant for the air sacs of the drosophila tracheal system. Dev Cell 3, 195–207 (2002).

4. Cabernard, C. & Affolter, M. Distinct roles for two receptor tyrosine kinases in epithelial branching morphogenesis in Drosophila. Dev Cell 9, 831–842 (2005).

5. Weaver, M. & Krasnow, M.A. Dual origin of tissue-specific progenitor cells in Drosophila tracheal remodeling. Science 321, 1496–1499 (2008).

6. Guha, A., Lin, L. & Kornberg, T.B. Organ renewal and cell divisions by differentiated cells in *Drosophila*. Proceedings of the National Academy of Sciences 105, 10832–10836 (2008).

7. Sato, M., Kitada, Y. & Tabata, T. Larval cells become imaginal cells under the control of homothorax prior to metamorphosis in the Drosophila tracheal system. Developmental Biology 318, 247–257 (2008).

8. Pitsouli, C. & Perrimon, N. Embryonic multipotent progenitors remodel the *Drosophila* airways during metamorphosis. Development 137, 3615–3624 (2010).

9. Pitsouli, C. & Perrimon, N. The homeobox transcription factor cut coordinates patterning and growth during Drosophila airway remodeling. Sci Signal 6, 2003424 (2013).

10. Jarecki, J., Johnson, E. & Krasnow, M.A. Oxygen Regulation of Airway Branching in *Drosophila* Is Mediated by Branchless FGF. Cell 99, 211–220 (1999).

11. Centanin, L., et al. Cell Autonomy of HIF Effects in Drosophila: Tracheal Cells Sense Hypoxia and Induce Terminal Branch Sprouting. Developmental Cell 14, 547–558 (2008).

12. Best, B.T. Single-cell branching morphogenesis in the Drosophila trachea. Dev Biol 451, 5–15 (2019).

13. Eilken, H.M. & Adams, R.H. Dynamics of endothelial cell behavior in sprouting angiogenesis. Curr Opin Cell Biol 22, 617–625 (2010).

14. Centanin, L., Gorr, T.A. & Wappner, P. Tracheal remodelling in response to hypoxia. J Insect Physiol 56, 447–454 (2010).

15. Gorr, T.A., Gassmann, M. & Wappner, P. Sensing and responding to hypoxia via HIF in model invertebrates. J Insect Physiol 52, 349–364 (2006).

16. Irisarri, M., et al. Central role of the oxygen-dependent degradation domain of Drosophila HIFalpha/Sima in oxygen-dependent nuclear export. Mol Biol Cell 20, 3878–3887 (2009).

17. Apidianakis, Y., Tamamouna, V., Teloni, S. & Pitsouli, C. Intestinal Stem Cells: A Decade of Intensive Research in Drosophila and the Road Ahead. (2017).

18. Jiang, H., Tian, A. & Jiang, J. Intestinal stem cell response to injury: lessons from Drosophila. Cell Mol Life Sci 73, 3337–3349 (2016).

19. Lemaitre, B. & Miguel-Aliaga, I. The digestive tract of Drosophila melanogaster. Annu Rev Genet 47, 377–404 (2013).

20. Jasper, H. Intestinal Stem Cell Aging: Origins and Interventions. Annu Rev Physiol 82, 203–226 (2020).

21. Micchelli, C.A. & Perrimon, N. Evidence that stem cells reside in the adult Drosophila midgut epithelium. Nature 439, 475–479 (2006).

22. Ohlstein, B. & Spradling, A. The adult Drosophila posterior midgut is maintained by pluripotent stem cells. Nature 439, 470–474 (2006).

23. Zeng, X. & Hou, S.X. Enteroendocrine cells are generated from stem cells through a distinct progenitor in the adult Drosophila posterior midgut. Development 142, 644–653 (2015).

24. Lin, G., Xu, N. & Xi, R. Paracrine Wingless signalling controls self-renewal of Drosophila intestinal stem cells. Nature 455, 1119–1123 (2008).

25. Xu, N., et al. EGFR, Wingless and JAK/STAT signaling cooperatively maintain Drosophila intestinal stem cells. Developmental Biology 354, 31–43 (2011).

26. Li, Z., Zhang, Y., Han, L., Shi, L. & Lin, X. Trachea-Derived Dpp Controls Adult Midgut Homeostasis in Drosophila. Developmental Cell 24, 133–143 (2013).

27. Miguel-Aliaga, I., Jasper, H. & Lemaitre, B. Anatomy and Physiology of the Digestive Tract of Drosophila melanogaster. Genetics 210, 357–396 (2018).

28. Kux, K. & Pitsouli, C. Tissue communication in regenerative inflammatory signaling: lessons from the fly gut. Frontiers in Cellular and Infection Microbiology 4(2014).

29. Linneweber, G.A., et al. Neuronal control of metabolism through nutrient-dependent modulation of tracheal branching. Cell 156, 69–83 (2014).

30. Campbell, K., et al. Collective cell migration and metastases induced by an epithelial-to-mesenchymal transition in Drosophila intestinal tumors. Nat Commun 10, 2311 (2019).

31. Jiang, H., et al. Cytokine/Jak/Stat signaling mediates regeneration and homeostasis in the Drosophila midgut. Cell 137, 1343–1355 (2009).

32. Buchon, N., Broderick, N.A., Poidevin, M., Pradervand, S. & Lemaitre, B. Drosophila intestinal response to bacterial infection: activation of host defense and stem cell proliferation. Cell Host Microbe 5, 200–211 (2009).

33. Pitsouli, C., Apidianakis, Y. & Perrimon, N. Homeostasis in infected epithelia: stem cells take the lead. Cell Host Microbe 6, 301–307 (2009).

34. Amcheslavsky, A., Jiang, J. & Ip, Y.T. Tissue damage-induced intestinal stem cell division in Drosophila. Cell Stem Cell 4, 49–61 (2009).

35. Biteau, B., Hochmuth, C.E. & Jasper, H. JNK activity in somatic stem cells causes loss of tissue homeostasis in the aging Drosophila gut. Cell Stem Cell 3, 442–455 (2008).

36. Markstein, M., et al. Systematic screen of chemotherapeutics in *Drosophila* stem cell tumors. Proceedings of the National Academy of Sciences 111, 4530–4535 (2014).

37. Chatterjee, M. & Ip, Y.T. Pathogenic stimulation of intestinal stem cell response in Drosophila. J Cell Physiol 220, 664–671 (2009).

38. Staley, B.K. & Irvine, K.D. Hippo signaling in Drosophila: recent advances and insights. Dev Dyn 241, 3–15 (2012).

39. Shaw, R.L., et al. The Hippo pathway regulates intestinal stem cell proliferation during Drosophila adult midgut regeneration. Development 137, 4147–4158 (2010).

40. Karpowicz, P., Perez, J. & Perrimon, N. The Hippo tumor suppressor pathway regulates intestinal stem cell regeneration. Development 137, 4135–4145 (2010).

41. Jiang, H. & Edgar, B.A. Intestinal stem cells in the adult Drosophila midgut. Exp Cell Res 317, 2780–2788 (2011).

42. Cordero, J.B., Stefanatos, R.K., Myant, K., Vidal, M. & Sansom, O.J. Non-autonomous crosstalk between the Jak/Stat and Egfr pathways mediates *Apc1*-driven intestinal stem cell hyperplasia in the *Drosophila* adult midgut. Development 139, 4524–4535 (2012).

43. Houtz, P., et al. Hippo, TGF-beta, and Src-MAPK pathways regulate transcription of the upd3 cytokine in Drosophila enterocytes upon bacterial infection. PLoS Genet 13, e1007091 (2017).

44. Sauer, H., Wartenberg, M. & Hescheler, J. Reactive oxygen species as intracellular messengers during cell growth and differentiation. Cell Physiol Biochem 11, 173–186 (2001).

45. Santabarbara-Ruiz, P., et al. ROS-Induced JNK and p38 Signaling Is Required for Unpaired Cytokine Activation during Drosophila Regeneration. PLoS Genet 11, e1005595 (2015).

46. Patel, P.H., et al. Damage sensing by a Nox-Ask1-MKK3-p38 signaling pathway mediates regeneration in the adult Drosophila midgut. Nat Commun 10, 4365 (2019).

47. He, L., et al. Antioxidants Maintain Cellular Redox Homeostasis by Elimination of Reactive Oxygen Species. Cell Physiol Biochem 44, 532–553 (2017).

48. Rojo de la Vega, M., Chapman, E. & Zhang, D.D. NRF2 and the Hallmarks of Cancer. Cancer Cell 34, 21–43 (2018).

49. Suh, J.H., et al. Decline in transcriptional activity of Nrf2 causes age-related loss of glutathione synthesis, which is reversible with lipoic acid. Proc Natl Acad Sci U S A 101, 3381–3386 (2004).

50. Brand, A.H. & Perrimon, N. Targeted gene expression as a means of altering cell fates and generating dominant phenotypes. Development 118, 401–415 (1993).

51. Shiga, Y., Tanaka-Matakatsu, M. & Hayashi, S. A nuclear GFP/β-galactosidase fusion protein as a marker for morphogenesis in living Drosophila. Development, Growth & Differentiation 38, 99–106 (1996).

52. Potter, C.J., Tasic, B., Russler, E.V., Liang, L. & Luo, L. The Q system: a repressible binary system for transgene expression, lineage tracing, and mosaic analysis. Cell 141, 536–548 (2010).

53. Sutherland, D., Samakovlis, C. & Krasnow, M.A. branchless encodes a Drosophila FGF homolog that controls tracheal cell migration and the pattern of branching. Cell 87, 1091–1101 (1996).

54. Grifoni, D., Sollazzo, M., Fontana, E., Froldi, F. & Pession, A. Multiple strategies of oxygen supply in Drosophila malignancies identify tracheogenesis as a novel cancer hallmark. Sci Rep 5, 9061 (2015).

55. Wang, C.W., Purkayastha, A., Jones, K.T., Thaker, S.K. & Banerjee, U. In vivo genetic dissection of tumor growth and the Warburg effect. Elife 5(2016).

56. Kaelin, W.G., Jr. The von Hippel-Lindau protein, HIF hydroxylation, and oxygen sensing. Biochem Biophys Res Commun 338, 627–638 (2005).

57. Klimova, T. & Chandel, N.S. Mitochondrial complex III regulates hypoxic activation of HIF. Cell Death Differ 15, 660–666 (2008).

58. Semenza, G.L. Hypoxia-inducible factors: coupling glucose metabolism and redox regulation with induction of the breast cancer stem cell phenotype. The EMBO Journal 36, 252–259 (2017).

59. Chandel, N.S., et al. Reactive oxygen species generated at mitochondrial complex III stabilize hypoxia-inducible factor-1alpha during hypoxia: a mechanism of O2 sensing. J Biol Chem 275, 25130–25138 (2000).

60. Brunelle, J.K., et al. Oxygen sensing requires mitochondrial ROS but not oxidative phosphorylation. Cell Metabolism 1, 409–414 (2005).

61. Guzy, R.D., et al. Mitochondrial complex III is required for hypoxia-induced ROS production and cellular oxygen sensing. Cell Metab 1, 401–408 (2005).

62. Movafagh, S., Crook, S. & Vo, K. Regulation of Hypoxia-Inducible Factor-1a by Reactive Oxygen Species: New Developments in an Old Debate. Journal of Cellular Biochemistry 116, 696–703 (2015).

63. Lavista-Llanos, S., et al. Control of the Hypoxic Response in *Drosophila melanogaster* by the Basic Helix-Loop-Helix PAS Protein Similar. Molecular and Cellular Biology 22, 6842–6853 (2002).

64. Centanin, L., Ratcliffe, P.J. & Wappner, P. Reversion of lethality and growth defects in Fatiga oxygen-sensor mutant flies by loss of hypoxia-inducible factor-alpha/Sima. EMBO reports 6, 1070–1075 (2005).

65. Majmundar, A.J., Wong, W.J. & Simon, M.C. Hypoxia-inducible factors and the response to hypoxic stress. Mol Cell 40, 294–309 (2010).

66. Tamamouna, V. & Pitsouli, C. The Hypoxia-Inducible Factor-1α in Angiogenesis and Cancer: Insights from the Drosophila Model. Gene Expression and Regulation in Mammalian Cells - Transcription Toward the Establishment of Novel Therapeutics, IntechOpen 72318, 209–241 (2018).

67. Deziel, E., et al. The contribution of MvfR to Pseudomonas aeruginosa pathogenesis and quorum sensing circuitry regulation: multiple quorum sensing-regulated genes are modulated without affecting lasRI, rhlRI or the production of N-acyl-L-homoserine lactones. Mol Microbiol 55, 998–1014 (2005).

68. Xiao, G., et al. MvfR, a key Pseudomonas aeruginosa pathogenicity LTTR-class regulatory protein, has dual ligands. Mol Microbiol 62, 1689–1699 (2006).

69. Liberati, N.T., et al. An ordered, nonredundant library of Pseudomonas aeruginosa strain PA14 transposon insertion mutants. Proc Natl Acad Sci U S A 103, 2833–2838 (2006).

70. Lee, K.A., et al. Bacterial-derived uracil as a modulator of mucosal immunity and gut-microbe homeostasis in Drosophila. Cell 153, 797–811 (2013).

71. Hochmuth, C.E., Biteau, B., Bohmann, D. & Jasper, H. Redox Regulation by Keap1 and Nrf2 Controls Intestinal Stem Cell Proliferation in Drosophila. Cell Stem Cell 8, 188–199 (2011).

72. Hanahan, D. & Weinberg, R.A. Hallmarks of cancer: the next generation. Cell 144, 646–674 (2011).

73. Zhai, Z., et al. Accumulation of differentiating intestinal stem cell progenies drives tumorigenesis. Nat Commun 6, 10219 (2015).

74. McGuire, S.E., Mao, Z. & Davis, R.L. Spatiotemporal gene expression targeting with the TARGET and gene-switch systems in Drosophila. Sci STKE 2004, pl6 (2004).

75. Buchon, N., et al. Morphological and molecular characterization of adult midgut compartmentalization in Drosophila. Cell Rep 3, 1725–1738 (2013).

76. Marianes, A. & Spradling, A.C. Physiological and stem cell compartmentalization within the Drosophila midgut. Elife 2, e00886 (2013).

77. Potente, M., Gerhardt, H. & Carmeliet, P. Basic and therapeutic aspects of angiogenesis. Cell 146, 873–887 (2011).

78. Trinchieri, G. Cancer and inflammation: an old intuition with rapidly evolving new concepts. Annu Rev Immunol 30, 677–706 (2012).

79. Folkman, J. Tumor angiogenesis: therapeutic implications. N Engl J Med 285, 1182–1186 (1971).

80. Folkman, J. Fundamental concepts of the angiogenic process. Curr Mol Med 3, 643–651 (2003).

81. Weis, S.M. & Cheresh, D.A. Tumor angiogenesis: molecular pathways and therapeutic targets. Nat Med 17, 1359–1370 (2011).

82. Choudhry, H. & Harris, A.L. Advances in Hypoxia-Inducible Factor Biology. Cell Metab 27, 281–298 (2018).

83. Lee, P., Chandel, N.S. & Simon, M.C. Cellular adaptation to hypoxia through hypoxia inducible factors and beyond. Nat Rev Mol Cell Biol 6, 020–0227 (2020).

84. Takenaga, K. Angiogenic signaling aberrantly induced by tumor hypoxia. Front Biosci (Landmark Ed) 16, 31–48 (2011).

85. Ohshiro, T. & Saigo, K. Transcriptional regulation of breathless FGF receptor gene by binding of TRACHEALESS/dARNT heterodimers to three central midline elements in Drosophila developing trachea. Development 124, 3975–3986 (1997).

86. André-Lévigne, D., Modarressi, A., Pepper, M.S. & Pittet-Cuénod, B. Reactive Oxygen Species and NOX Enzymes Are Emerging as Key Players in Cutaneous Wound Repair. Int J Mol Sci 18(2017).

87. Jia, Y.-T., et al. Activation of p38 MAPK by Reactive Oxygen Species Is Essential in a Rat Model of Stress-Induced Gastric Mucosal Injury. The Journal of Immunology 179, 7808–7819 (2007).

88. Warren, C.M., Ziyad, S., Briot, A., Der, A. & Iruela-Arispe, M.L. A ligand-independent VEGFR2 signaling pathway limits angiogenic responses in diabetes. Science signaling 7, ra1-ra1 (2014).

89. Nezu, M., et al. Nrf2 inactivation enhances placental angiogenesis in a preeclampsia mouse model and improves maternal and fetal outcomes. Sci Signal 10(2017).

90. Reczek, C. & Chandel, N. The Two Faces of Reactive Oxygen Species in Cancer. Annual Review of Cancer Biology 1(2017).

91. Perez, E., Lindblad, J.L. & Bergmann, A. Tumor-promoting function of apoptotic caspases by an amplification loop involving ROS, macrophages and JNK in Drosophila. Elife 6(2017).

92. Ha, E.-M., et al. An Antioxidant System Required for Host Protection against Gut Infection in *Drosophila*. Developmental Cell 8, 125–132 (2005).

93. Lee, W.J. & Brey, P.T. How microbiomes influence metazoan development: insights from history and Drosophila modeling of gut-microbe interactions. Annu Rev Cell Dev Biol 29, 571–592 (2013).

94. Jones, R.M., et al. Symbiotic lactobacilli stimulate gut epithelial proliferation via Nox-mediated generation of reactive oxygen species. The EMBO Journal 32, 3017–3028 (2013).

95. Kim, S.H. & Lee, W.J. Role of DUOX in gut inflammation: lessons from Drosophila model of gut-microbiota interactions. Front Cell Infect Microbiol 3, 116 (2014).

96. Gervais, L. & Casanova, J. The *Drosophila* homologue of SRF acts as a boosting mechanism to sustain FGF-induced terminal branching in the tracheal system. Development 138, 1269–1274 (2011).

97. Bardin, A.J., Perdigoto, C.N., Southall, T.D., Brand, A.H. & Schweisguth, F. Transcriptional control of stem cell maintenance in the Drosophila intestine. Development 137, 705–714 (2010).

98. Kumar, J.P. & Moses, K. EGF receptor and Notch signaling act upstream of Eyeless/Pax6 to control eye specification. Cell 104, 687–697 (2001).

99. Ha, E.M., Oh, C.T., Bae, Y.S. & Lee, W.J. A direct role for dual oxidase in Drosophila gut immunity. Science 310, 847–850 (2005).

100. Thibault, S.T., et al. A complementary transposon tool kit for Drosophila melanogaster using P and piggyBac. Nature Genetics 36, 283–287 (2004).

101. Lee, T. & Luo, L. Mosaic analysis with a repressible cell marker (MARCM) for Drosophila neural development. Trends Neurosci 24, 251–254 (2001).

102. Pitsouli, C. & Delidakis, C. The interplay between DSL proteins and ubiquitin ligases in Notch signaling. Development 132, 4041–4050 (2005).

103. Geiszt, M. & Leto, T.L. The Nox family of NAD(P)H oxidases: host defense and beyond. J Biol Chem 279, 51715–51718 (2004).

104. Bonilla, E., Medina-Leendertz, S., Villalobos, V., Molero, L. & Bohorquez, A. Paraquat-induced oxidative stress in drosophila melanogaster: effects of melatonin, glutathione, serotonin, minocycline, lipoic acid and ascorbic acid. Neurochem Res 31, 1425–1432 (2006).

105. Vrailas-Mortimer, A., Gomez, R., Dowse, H. & Sanyal, S. A survey of the protective effects of some commercially available antioxidant supplements in genetically and chemically induced models of oxidative stress in Drosophila melanogaster. Exp Gerontol 47, 712–722 (2012).

